# Augmented Transcutaneous Stimulation Using an Injectable Electrode

**DOI:** 10.1101/2021.09.29.462482

**Authors:** Nishant Verma, Robert D Graham, Jonah Mudge, James K Trevathan, Manfred Franke, Andrew J Shoffstall, Justin Williams, Ashley N Dalrymple, Lee E Fisher, Doug J Weber, Scott F Lempka, Kip A Ludwig

## Abstract

Minimally invasive neuromodulation technologies seek to marry the neural selectivity of implantable devices with the low-cost and non-invasive nature of transcutaneous electrical stimulation (TES). The Injectrode^®^ is a needle-delivered electrode that is injected onto neural structures under image guidance. Power is then transcutaneously delivered to the Injectrode using surface electrodes. The Injectrode serves as a low-impedance conduit to guide current to the deep on-target nerve, reducing activation thresholds by an order of magnitude compared to using only surface stimulation electrodes. To minimize off-target recruitment of cutaneous fibers, the energy transfer efficiency from the surface electrodes to the Injectrode must be optimized.

TES energy is transferred to the Injectrode through both capacitive and resistive mechanisms. Electrostatic finite element models generally used in TES research consider only the resistive means of energy transfer by defining tissue conductivities. Here, we present an electroquasistatic model, taking into consideration both the conductivity and permittivity of tissue, to understand transcutaneous power delivery to the Injectrode. The model was validated with measurements taken from (n=4) swine cadavers. We used the validated model to investigate system and anatomic parameters that influence the coupling efficiency of the Injectrode energy delivery system. Our work suggests the relevance of electroquasistatic models to account for capacitive charge transfer mechanisms when studying TES, particularly when high-frequency voltage components are present, such as those used for voltage-controlled pulses and sinusoidal nerve blocks.

## Introduction

Non-invasive transcutaneous electrical stimulation (TES) therapies seek to directly manipulate neural activity to ameliorate disease. Because they are non-invasive, TES devices are generally low-cost and low-risk. While these completely non-invasive devices can engage deep nerves, they also activate more superficial neural structures, such as cutaneous receptors in the skin and off-target superficial nerves (Bucksot et al., 2020). Activation of superficial off-target neural structures leads to side effects, including noxious sensation and uncomfortable muscular contractions (Bucksot et al., 2020; Manson et al., 2020), which limit the stimulation dose from being increased to engage the deep on-target nerve.

An implantable device that directly interfaces with the nerve can achieve on-target neural engagement in a more specific manner (Aristovich et al., 2021). However, traditional implantable devices consisting of an implanted battery, electronics, leads, and stimulation electrodes are complex and costly (Kumar and Bishop, 2009; Udo et al., 2012). The complexity of the implanted device and the associated supply chain, in which manufacturing procedures must be tightly controlled across several suppliers, contributes to the cost of the therapy and is prone to multiple points of failure (FDA, 1997; Carome, 2020). Once manufactured, a traditional implantable device requires invasive surgical placement, which adds to the cost of the therapy (Trevathan, 2019; Kumar and Bishop, 2009). Traditional implanted devices also use pre-formed, rigid neural interfacing electrodes that do not conform well to complex neural structures (He et al., 2020).

Minimally invasive neuromodulation therapies are designed to attain the spatial specificity and neural target engagement depth of an implantable therapy while maintaining the accessibility and low-risk attributes of non-invasive TES. Examples of current minimally invasive technologies are found in Supplementary Material 1. These minimally invasive systems generally consist of a small device implanted at the target neural structure and an external power source, which powers the implant (Loeb et al., 2006; Ilfeld et al., 2021). However, these devices are still complex and costly and use stiff neural interfacing electrodes that do not conform to the target neural structure to reduce tissue trauma and better isolate the target nerve (Loeb et al., 2006).

The Injectrode is a minimally invasive neuromodulation electrode technology designed to provide the selectivity of an implanted electrode with a more favorable risk profile (Trevathan et al., 2019; Dalrymple et al., 2021). It can be injected using a syringe and forms in-body onto the neural structure to better isolate the target nerve, even with individual anatomical differences. The pre-curing flexibility of the Injectrode allows it to conform around complex neural structures, including nerve plexi found close to target end-organs, which is difficult with conventional pre-formed neurostimulation electrodes. Therefore, the Injectrode may extend the range of nerves that can be targeted with neurostimulation and improve end-organ targeting specificity. Additional Injectrode material is extruded as part of the injection to form a conductive conduit connecting the deep nerve target to just under the surface of the skin. Finally, more Injectrode is injected just under the skin to create a disc shaped ‘collector’. This collector couples with external non-invasive TES electrodes to transfer charge delivered from a non-invasive TES unit and route it to the deep target nerve. Fig. 1 (a) illustrates the delivery procedure of the Injectrode system.

**Figure 1:**
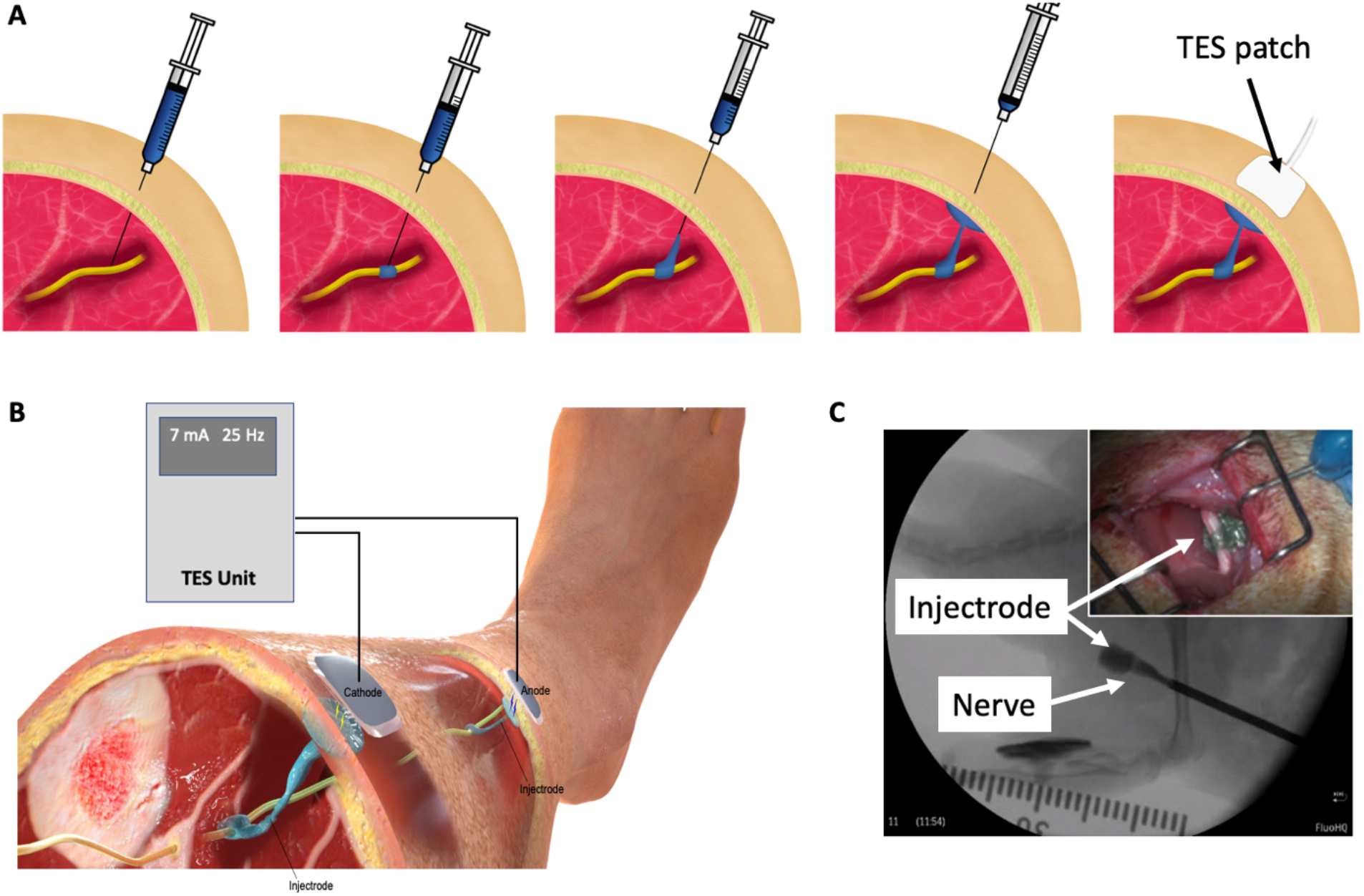
**(a)** Delivery procedure of the Injectrode system. The Injectrode is injected onto a neural structure. A syringe containing the Injectrode is deployed to the target nerve under image guidance. The Injectrode is deployed to form an interface with the nerve. The syringe is then drawn back while injecting the Injectrode – to form a conductive path from the deep nerve to skin. More Injectrode material is then injected under the skin to form a ‘collector’. An externally placed TES patch electrode non-invasively delivers charge to the Injectrode. **(b)** Injectrode system in bipolar configuration after deployment. A TES unit is used to deliver energy non-invasively to the Injectrode collectors. The Injectrode sets up a low-impedance conduit to guide current to the deep target nerve. **(c)** Injectrode delivery onto a neural structure under image guidance. Opacity in the figure corresponds to the Injectrode’s thickness with a portion going around the nerve showing lightest opacity. (Inset top right) Injectrode conforming to neural structure.

Energy is delivered to the Injectrode by external surface electrodes using low-frequency electric fields through both capacitive and resistive mechanisms. Capacitively, the external surface electrode and the in-body subcutaneous collector act like two plates of a capacitor with skin as the dielectric. High-frequency components of the electric field are transferred preferentially through this capacitive route. Simultaneously, current is also transferred through a resistive route. The surface electrodes set up an electric potential gradient in tissue, visualized in Fig. 2 (d). The Injectrode collector is placed subcutaneously, close to the surface electrodes, and provides a low-impedance conduit for current to flow from one collector, down the Injectrode lead, through the nerve and other tissue, and back up the other Injectrode lead and collector. If the path formed by the Injectrode is of lower impedance than a direct path between the two collectors through tissue, current preferentially travels through the Injectrode path, stimulating the deep on-target nerve in the process. The concept of power transfer across the skin from surface electrodes to implanted electrodes using low-frequency electric fields has been previously presented (Gan and Prochazka, 2007; Gan and Prochazka, 2010; Gaunt and Prochazka, 2009), with the Injectrode system now providing a minimally invasive injectable implementation of the concept.

**Figure 2:**
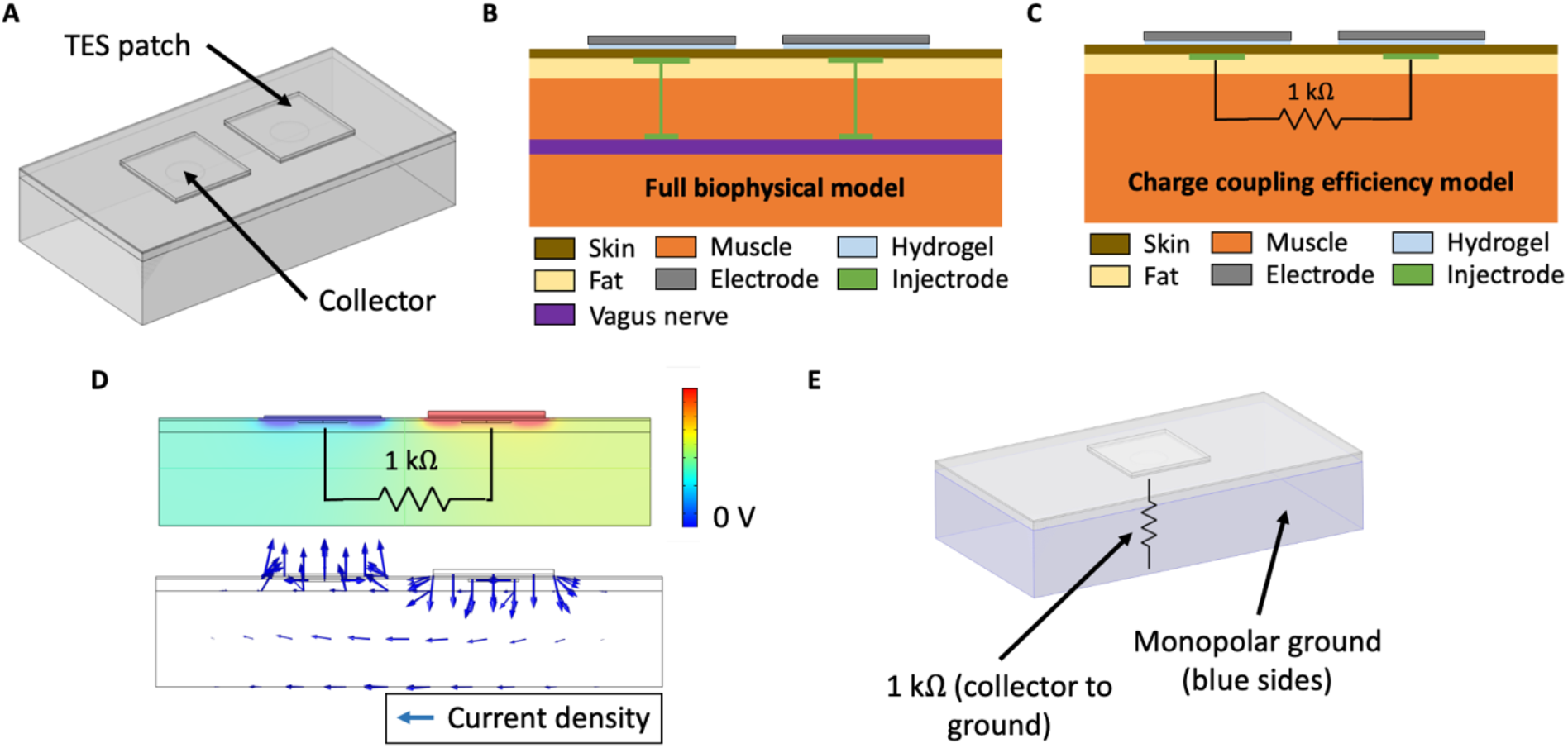
**(a)** Three-layer tissue model of the Injectrode system in COMSOL Multiphysics to study transcutaneous charge transfer from the surface electrodes to the subcutaneous collectors. **(b)** Schematic of the Injectrode system FEM model. This full biophysical model was used to study on- and off-target neural recruitment with the Injectrode system. **(c)** This charge coupling efficiency model was used to study the transcutaneous charge transfer from the surface electrodes to the subcutaneous collectors. The 1 kΩ resistor was used to represent the impedance of the Injectrode connection to the deep nerve, the Injectrode-nerve interface, the nerve, and the leakage between the two Injectrode conduction paths. **(d)** (Top) Electric potential solution for the standard model configuration at 10 kHz. Electric potential difference between the two subcutaneous collectors causes current to flow through the 1 kΩ resistor connecting the two collectors. (Bottom) Arrows representing current density flow (1 kΩ resistor not shown). **(e)** Monopolar configuration of the Injectrode system to study the transcutaneous power transfer from the surface electrode to the subcutaneous Injectrode collector. Here, the 1 kΩ resistor is connected from the single collector to a 0 V ground potential.

The Injectrode system is a platform technology that could be applied to various neuromodulation indications. To minimize off-target recruitment of cutaneous and superficial fibers by the surface electrodes, energy transfer efficiency from the surface electrodes to the Injectrode must be optimized. Investigating power transfer across the skin to the Injectrode collector requires consideration of charge transfer due to both resistive and capacitive means. Finite element analysis in the TES field traditionally considers charge transfer only due to resistive means using an electrostatic analysis (Kuhn et al., 2009). An electrostatic, or direct-current (DC), analysis ignores the capacitive displacement current that accompanies oscillating electric fields, which is essential to understand the transcutaneous capacitive charge coupling behavior of the Injectrode. Previous work has established the importance of dielectric properties of tissue and capacitive charge transfer even when studying TES therapies (Medina and Grill, 2014; Poulsen et al., 2020; Kuhn et al., 2009). The skin acts as a dielectric and allows direct coupling of higher-frequency components in the TES waveform from surface electrodes to the underlying tissue. The placement of subcutaneous Injectrode collectors increases this capacitive coupling and resultant capacitive charge transfer.

Here, we use the finite element method (FEM) to develop an electroquasistatic model for the Injectrode system to study power transfer from the surface electrodes to the Injectrode collector by both capacitive and resistive means. We selected model dimensions considering the neck region in humans. The desired on-target effect was recruitment of the vagus nerve and undesired off-target effect was activation of cutaneous and superficial fibers responsible for paresthesia or lip curl (likely due to activation of the cervical branch of the facial nerve innervating the platysma muscle, as seen in use of the gammaCore non-invasive vagus nerve stimulation (nVNS) device (Nonis et al., 2017)). We validated the model output in swine cadavers and then used the validated model to investigate Injectrode system and anatomic parameters (e.g., tissue thickness and conductivity) that influence the coupling efficiency of charge delivery to the Injectrode collector. Finally, we performed biophysical modeling to investigate how the Injectrode alters recruitment of on- and off-target neural structures in comparison to traditional TES. These results provide insights into waveform design and system parameters for the optimization of the Injectrode system to achieve on-target activation of deep neural structures while minimizing off-target activation of superficial neural fibers.

## Methods

### Electroquasistatic FEM model

COMSOL Multiphysics version 5.5 software (COMSOL, Burlington, MA) was used to create and solve the FEM model for electric field and currents. A three-dimensional model was set up using the Electric Currents physics interface under the AC/DC module. The Electric Currents physics interface computes both ohmic (resistive) and displacement (capacitive) currents by considering tissue conductivity and permittivity, respectively, while ignoring inductive effects (Bossetti et al., 2008). A three-layer tissue model consisting of skin, fat, and muscle was set-up as shown in Fig. 2 (a). The Injectrode system was constructed using two square surface electrodes and two subcutaneous circular collector electrodes. To isolate and investigate the effects of system parameters on transcutaneous coupling between the surface electrodes and collectors, we used a defined electrical load between the two collectors (Fig. 2 (c)) implemented in the Electric Circuit physics interface in COMSOL. The electrical load was defined as a 1 kΩ resistor, based on electrical impedance spectroscopy (EIS) measurements from Trevathan et al. (2019), to model the impedance of the Injectrode connection from the collector down to the nerve, the Injectrode-nerve interface, the nerve impedance, and the leakage between the two Injectrode conduction paths (Fig. 2 (b)). Boundary conditions for the external tissue surfaces were set to zero normal current and an initial condition of 0 V. In the monopolar configuration, shown in Fig. 2 (e), ground was set as the five surfaces (left, right, bottom, front, and back) of the muscle layer and an end of the 1 kΩ resistor was set to the 0 V ground potential. The other external tissue surfaces in the monopolar model were set to the same boundary conditions as in the bipolar model. The model was run with stationary (DC), time-dependent (transient), and frequency-domain studies. The frequency-domain study used a complex analysis denoting ohmic currents as real and displacement currents as imaginary. All currents and voltages reported are absolute values.

Internal current (Gan and Prochazka, 2010), or nerve current, was measured in the model as the current flowing through the 1 kΩ resistor and denoted as I_Nerve_ (mA). External current or TES current was measured as the current delivered by the external TES electrode and denoted as I_TES_ (mA), with the corresponding voltage to drive the TES current denoted as V_TES_ (V). The simplified three-layer model allowed study of the transcutaneous coupling behavior of the surface electrodes with the subcutaneous Injectrode collectors. A capture ratio (Gan and Prochazka, 2010) of I_Nerve_/I_TES_ was calculated to estimate the efficiency of the transcutaneous current delivery to current arriving at the modeled deep nerve and reported as ‘efficiency’ (%). Lastly, surface electrode current density was used as a proxy for off-target cutaneous fiber activation (Slopsema et al., 2018) and I_Nerve_ was used as a proxy for on-target neural fiber activation.

Model geometry was set to replicate the parameters of a 50-60 kg domestic swine’s lower abdomen, on which the model was validated, and is similar to human tissue thicknesses at the neck. The human neck represents a possible target to access the vagus nerve for an Injectrode deployment. Tissue thickness was set as 1 mm for skin – typical of the measurements made in the swine model at the abdomen (mean=1.01 mm; standard deviation (SD)=0.31 mm; n=16 measurements). However, the skin at the neck is thicker in swine. A skin thickness of 1 mm is also representative of the mean thickness of human skin at the neck (mean=1.3 mm; SD=0.2 mm) (Hoffmann et al., 1994). Fat and muscle thickness were set at 5 mm and 40 mm, respectively, to represent the measured values in swine. These values are representative of the human neck region (Störchle et al., 2018). A FEM model area of 21 cm by 11 cm was studied, leaving a minimum border of 2 cm between the surface electrodes and model edges to reduce interference from edge effects. The model size was varied to ensure that a larger area would not results in a difference of more than 1% in nerve current (I_Nerve_) (Poulsen et al., 2020).

Several steps were taken to ensure proper selection of mesh size and time steps for the FEM analysis. ‘Physics’ settings in the COMSOL software were used to efficiently define the mesh properties and time steps, allowing for shorter intervals in regions of greater parameter gradients. Mesh density was progressively made coarser to decrease computation time while nerve current (I_Nerve_) remained within 1% of the finest mesh size (Kuhn et al., 2009; Poulsen et al., 2020). The direct solver was used to solve the stiffness matrix. Computations were run locally on a Windows 10 desktop with a 3.00 GHz Intel i7 processor and 32 GB of RAM.

### Tissue conductivity and permittivity values

The selection of appropriate skin, fat, and muscle conductivity and permittivity values is important to construct an accurate model. We used values from the well-established Gabriel et al. (1996b) database – summarized in Table 1. All values were drawn from the same database to prevent bias that may arise when selecting values from multiple sources. In COMSOL, the “contact impedance” was set to 6.9 x 10^-2 Ω.m^2^, based on empirical measurements detailed in Supplementary Material 2, to define all Injectrode-tissue electrochemical interfaces.

**Table 1:** Material electrical properties used in the FEM model

| Tissue | Conductivity (S/m) | Relative permittivity | Source |
| --- | --- | --- | --- |
| Skin | $1.80 \times 10^{-4}$ | $1.17 \times 10^3$ | Human, 37 °C, 1 kHz, dry (Gabriel et al., 1996b) |
| Fat | $2.46 \times 10^{-2}$ | $2.08 \times 10^4$ | Bovine, 37 °C, 1 kHz, non-infiltrated fat (Gabriel et al., 1996b) |
| Muscle | $5.23 \times 10^{-1}$ | $1.24 \times 10^6$ | Ovine, 37 °C, 1 kHz, parallel muscle fibers (Gabriel et al., 1996b) |
| Hydrogel | $1.6 \times 10^{-2}$ | $1.4 \times 10^6$ | Measured, see Supplementary Material 2 |
| Injectrode | $3.774 \times 10^7$ | 1 | COMSOL in-built value for a conductive metal |

Dispersive properties, or frequency-dependent conductivity and permittivity, were not considered in the current model. Literature values (Gabriel et al., 1996b) show that skin, fat, and muscle conductivity and skin permittivity do not change substantially within the 10 Hz – 25 kHz frequency range, which is the range relevant to this study. Fat and muscle permittivity values are a function of frequency in this range but were considered a second-order effect in this simplified model. Frequency-dependent conductivity and permittivity may be added to the model – improving its accuracy (Zander et al., 2020) but in exchange for increased mathematical complexity and computation time.

### Cadaver validation of FEM model

To validate the FEM model, measurements were taken from the abdominal region of domestic swine within two hours of death. Tissue has been reported to preserve its electrical properties for a few minutes to hours after death (Foster and Schwan, 1989). Skin at the abdominal region of the pig is similar in thickness to human skin at the neck (Hoffmann et al., 1994). The domestic swine is a good model for human skin, with similar histological and biochemical properties, such as epidermal turnover time, subdermal adipose tissue, vasculature, and collagen structure, and arrangement in the dermis (Avci et al., 2013). Although domestic swine do not possess eccrine sweat glands, they have apocrine glands distributed throughout their skin surface (Avci et al., 2013). The protocol described below was developed through (n=5) preliminary cadaver experiments (data not shown). Once the final protocol was established, confirmatory measurements were taken from both sides of (n=4) swine cadavers and are presented here.

An approximately 2 cm incision was made in the skin using a scalpel. Forceps were used to bluntly dissect a space between the skin and fat layer, creating a pocket for the collector to be inserted. The creation of the incision a distance from the surface electrodes prevented current from routing through a break in the skin. Similar to Gan and colleagues (Gan and Prochazka, 2010), a stainless-steel disk (2.1 cm diameter) was used as a consistent representation of an Injectrode collector for comparison to FEM model outputs, because an Injectrode collector varies from deployment to deployment due to local tissue consistency and conformance. Fig. 4 (a) shows the similar size and shape between a stainless-steel disc and the cured Injectrode collector. This similar size and shape ensures similar results for V_TES_, I_TES_, and I_Nerve_. A second 2 cm incision was made to insert the second collector. A 1 kΩ resistor was connected externally between the two collectors (between the two red wires running out of the incision on the left side of Fig. 4 (a)), which allowed us to precisely compare cadaver data to the paired FEM model. Surface electrodes 5 x 5 cm in size (TENSpros, Saint Louis, MO) were applied with an edge-to-edge separation of 2 cm. Prior to taking measurements, 5 minutes were allowed for the surface electrodes to equilibrate with skin.

After equilibration, stability of the electrode-skin interface was verified by delivering ten 300 *μ*s current-controlled pulses at 19 mA and measuring the voltage across the TES electrodes. Next, our test stimulation waveforms were delivered at the surface TES electrodes using an AM 4100 isolated high-power stimulator (AM Systems, Sequim, WA). Stimulus was delivered at 19 mA for current-controlled and 28 V for voltage-controlled waveforms, below the voltage at which electroporation occurred. Electroporation, marked by a sudden decrease in electrode-skin impedance, was observed during preliminary testing at the swine neck when a stimulus over 90 V was applied during a 300 *μ*s pulse. Several waveforms, including monophasic pulses of varying rise times, were tested. Waveform test order was randomized. The initial 300 *μ*s current-controlled pulse at 19 mA was delivered again at the end of testing to check for degradation over time in the cadaver model. All measurements were made within 20 minutes of incision to reduce fluid buildup in the surgical pockets. No appreciable degradation was measured during any confirmatory experiment. The procedure was then repeated on the contralateral side of the abdomen.

Nerve current (I_Nerve_), i.e., internal current (Gan and Prochazka, 2010), was calculated from voltage measurements taken across the 1 kΩ resistor connected between the two collectors using a TMDP0200 high-voltage differential probe (Tektronix, Beaverton, OR). V_TES_, the voltage across the TES electrodes, was measured using an identical differential probe. Both differential probes were connected to a DPO2004B oscilloscope (Tektronix, Beaverton, OR) with an external probe power supply. A Keithley DAQ 6510 (Tektronix, Beaverton, OR) measured the current to the TES electrode (I_TES_). It was crucial to use a setup that was well isolated from ground for the appropriate frequency range under measurement. The AM 4100 isolated stimulator was not well isolated from ground at higher-frequency voltages; therefore, in confirmatory experiments, differential probes were used to achieve electrical isolation. After waveform testing, the skin between the surface electrode and collector was cut with a scalpel and its thickness was measured using a Vernier caliper to be (mean=1.01 mm; SD=0.31 mm; n=16 measurements).

### Multi-compartment neuron model

To study on-target (i.e., vagus) and off-target (i.e., cutaneous) neuronal activation, we extended the validated FEM model used to study transcutaneous charge coupling (Fig. 3 (a)) and implemented multicompartment cable models of cutaneous and vagal axons using the NEURON simulation environment (v7.7) (Hines et al., 1997). Vagus nerve stimulation is believed to activate sensory fibers for its therapeutic benefits (Krahl, 2012). Aβ fibers are the lowest activation threshold sensory fibers in the cervical vagus and may feasibly be recruited during clinical stimulation (Nicolai et al. 2020; Krahl, 2012). We implemented previously described axon models of an Aβ-low-threshold mechanoreceptor (LTMR) in both the vagus and cutaneous regions (canonically responsible for paresthesia), and an Aδ-high-threshold mechanoreceptor (HTMR) in the cutaneous region (canonically responsible for noxious sensations). Briefly, we modeled each axon morphology using the double-cable McIntyre-Richardson-Grill (MRG) model of a myelinated mammalian peripheral axon (McIntyre et al., 2002). The MRG axon model is parametrized for discrete axonal diameters. Therefore, we modeled a 10.0 μm diameter Aβ-axon and a 2.0 μm diameter Aδ-axon to approximate the axonal diameters used in previous modeling studies of TES (Tigerholm et al., 2019). We modeled the membrane dynamics of each axon using previously described ion channel properties of Aβ-LTMRs (Graham et al., 2019) and Aδ-HTMRs (Graham et al., 2020), which reproduced experimental data describing action potential characteristics and conduction velocities found in sensory neurons.

**Figure 3:**
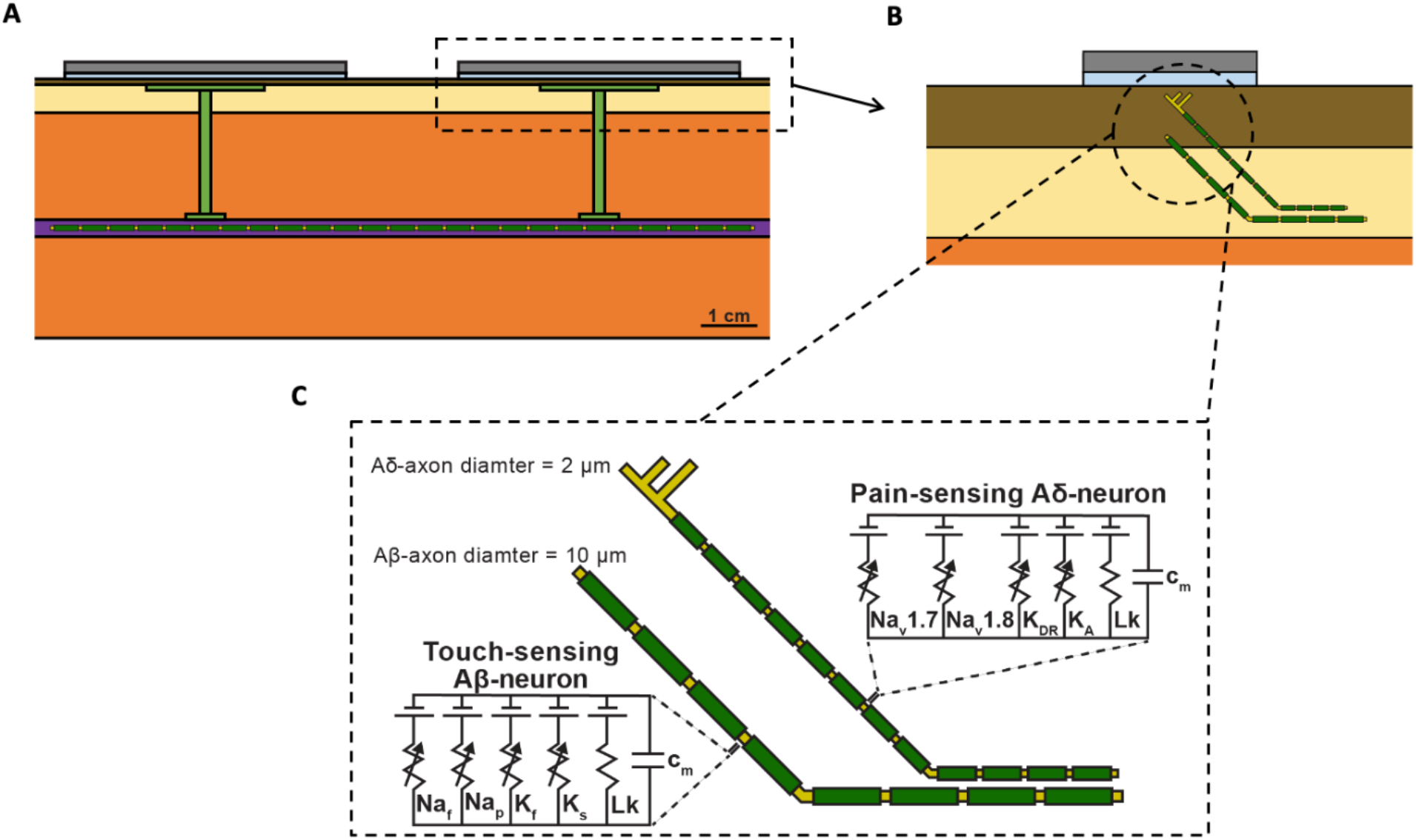
**(a)** Full FEM model used in the biophysical study. The 1 kΩ resistor between the two collectors in the simplified FEM model was replaced with the vagus nerve, Injectrode connections down to the vagus nerve, and Injectrode interfaces with the vagus nerve. The vagus nerve was populated with axons. **(b)** Zoomed view of dotted box in (a) showing the skin populated with cutaneous fibers. **(c)** Biophysical equivalent circuit model of cutaneous Aβ and Aδ neural fibers.

We then distributed each axon type throughout the bipolar Injectrode FEM model. A modified FEM model was used for the biophysical study, additionally incorporating the vagus nerve, Injectrode leads down to the vagus nerve, and Injectrode interfaces with the vagus nerve (Fig. 3 (a)) all of which were previously modeled as a 1 kΩ resistor between the two collectors (Fig. 2 (c)). The previous simplified FEM model allowed direct interpretation of the results as being caused by changes in the transcutaneous charge coupling efficiency between the surface electrodes and collectors. For the biophysical investigation, we created two populations of axon trajectories: a population of axons in the vagus nerve, and a population of cutaneous afferents that terminated below the active TES patch electrode (Fig. 3 (b)). To generate the vagus axon population, we created a two-dimensional regular grid parallel to the face of the vagus nerve with 100 μm spacings in all directions. Each point on the grid acted as a seed point for a vagus axon, which then traveled in a straight line to the other end of the nerve.

We also modeled Aβ- and Aδ-cutaneous axons, morphologically and anatomically similar to the cutaneous afferent models developed by Tigerholm and colleagues (Tigerholm et al., 2019). The cutaneous axon population was constructed using a two-dimensional regular grid parallel to the active TES patch with 1.5 mm spacings in all directions, 900 μm beneath the surface of the skin. This grid extended 5 mm beyond the edge of the TES patch. Each point on the grid acted as a terminal point for an Aβ-cutaneous axon, responsible for transmitting mechanosensations, which may result in paresthesias (Tigerholm et al., 2019). Cutaneous Aδ-axons terminate more superficially than Aβ-axons and are responsible for transmitting noxious sensations (Tigerholm et al., 2019; Mørch et al., 2011). Therefore, we generated a separate two-dimensional regular grid parallel to the active TES patch, 500 μm beneath the surface of the skin to serve as terminal points for cutaneous Aδ-axons. Each cutaneous axon traveled parallel to the vagus nerve 4 mm under the surface of the skin in the subcutaneous fat layer of the FEM model, before branching up towards the skin (Tigerholm et al., 2019). Cutaneous Aβ-axons terminated in a passive node of Ranvier, while cutaneous Aδ-axons terminated in a passive branching structure using a previously described morphology, which reproduces nerve fiber densities found in human skin (Tigerholm et al., 2019).

### Simulating the neural response to the Injectrode system

We interpolated the extracellular potentials calculated by the FEM model onto the middle of each neural compartment and used NEURON’s extracellular mechanism within the Python programming environment to simulate the axonal response to the Injectrode system (Hines et al., 2009). We used a backward Euler implicit integration method with a time step of 5 μs to calculate each compartment’s time-varying transmembrane voltage in response to Injectrode-TES stimulation (Graham et al., 2019). Our goal was to investigate how the Injectrode system affects the activation of on- and off-target axons. Therefore, we calculated each axon’s activation threshold, i.e., the minimum current amplitude needed to induce a propagating action potential, using a binary search algorithm with a resolution of 1 μA. For all biophysical simulations, we used a current-controlled stimulus pulse width of 300 μs.

## Results

This work establishes a FEM model of the Injectrode system to study transcutaneous charge coupling using low-frequency electric fields. An electroquasistatic model was set up in COMSOL, solving for both ohmic (resistive) and displacement (capacitive) current to study transcutaneous charge coupling in the Injectrode system. The transcutaneous coupling FEM model was a simplified model used to isolate changes due to coupling behavior between the surface electrodes and the collector. The transcutaneous coupling FEM model was validated with measurements of several waveforms on both the left and right side of recently dead swine (n=4). The validated model was used to investigate the Injectrode system and patient-dependent parameters (e.g., surface electrode placement, tissue thickness, skin preparation, tissue electrical properties) most sensitive to coupling efficiency (ratio between nerve current and externally applied surface electrode current). Maximizing the efficiency ratio minimizes surface electrode current, which activates off-target cutaneous and superficial nerves, while maximizing the current available at the deep on-target nerve. Finally, a full biophysical model was used to investigate on-versus off-target neural recruitment.

### Transcutaneous coupling FEM model output and cadaver validation

Fig. 4 (b-c) show simulation results (solid line) compared to cadaver measurements (shaded area representing ± 1 SD of n=8 measurements) used to validate the FEM model. Seen in Fig. 4 (b), the current drawn at the TES electrode patch (blue) with a symmetric trapezoidal voltage pulse (red) mimics the current seen at the deep nerve (green). As the TES-tissue interface is substantially capacitive, the current generated is more sensitive to the change in voltage applied dV_TES_/dt than absolute voltage (V_TES_) as seen in Fig. 4 (b-c). Supplementary Material 3 shows a similar figure for a current-controlled stimulation pulse.

**Figure 4:**
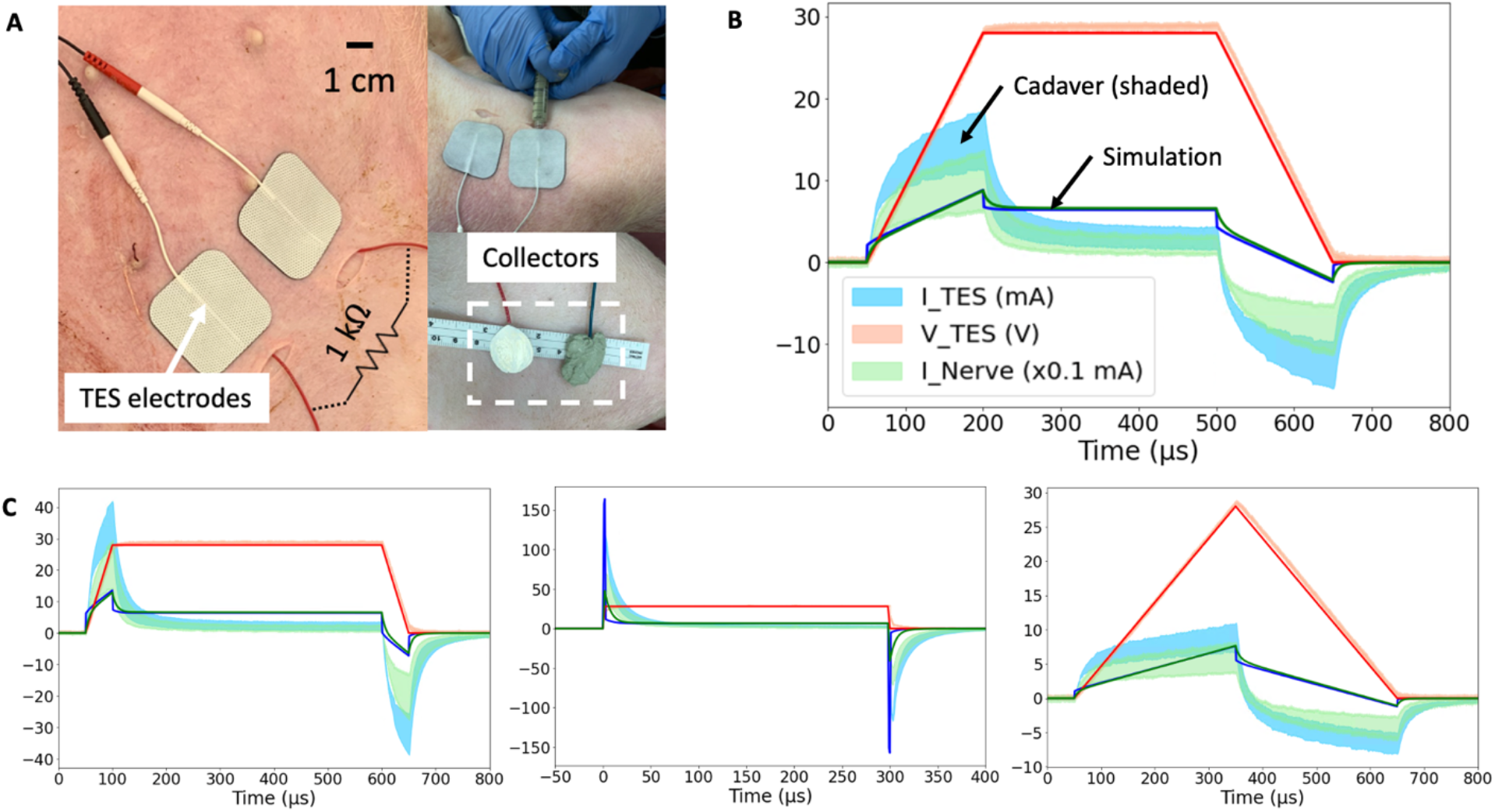
**(a)** Domestic swine cadaver verification of the FEM model using stainless-steel discs. **(b)** 28 V voltage-controlled 600 *μ*s pulses with 150 *μ*s rise and fall times. Three solid lines are simulation results, and three shaded areas are cadaver measurements ± 1 SD (n=8 measurements from both sides of n=4 cadavers). Red solid line (simulation) and shaded area (cadaver validation measurements) represent voltage of applied stimulation waveform, blue represents current through surface electrodes, and green represents nerve current (scaled by x0.1 mA for visualization). **(c)** 28 V voltage-controlled pulses of 600 *μ*s duration with 50 *μ*s rise and fall times (left), 300 *μ*s duration with fastest (~2 μs) rise and fall time (center), 600 *μ*s duration with 300 *μ*s rise and fall times (right). Note: 50 *μ*s rise time (left) is n=7 measurements due to the incorrect application of waveform amplitude in one sample.

A controlled rise time in surface electrode voltage could be used to exploit the additional displacement current provided by capacitive coupling without increasing the absolute voltage applied across the surface electrodes. The displacement current through a capacitor is proportional to the rate of change of voltage across it (dV/dt). Several rise times corresponding to dV_TES_/dt of 0.6, 0.2, and 0.1 V/μs were investigated in the FEM model and cadaver. Faster rise times corresponded to greater surface electrode current and nerve current as shown in Fig. 4 (b-c). In this manner, controlled rise times on voltage-controlled waveforms can be used to add additional displacement current to the ohmic nerve current.

General waveform shapes were well captured by the FEM model. For example, in Fig. 4 (c, left), the TES Voltage (V_TES_) represents the applied voltage-controlled waveform with a 50 *μ*s rise and fall time. During the rise and fall, the exponential charging shape of TES Current (I_TES_) means the TES electrodes are capacitively coupling with tissue and the collectors, and displacement current dominates. The identical shapes of I_TES_ and the nerve current (I_Nerve_) means that once current enters tissue, ohmic current transfer dominates.

Differences in absolute values between the model and cadaver measurements can be explained by differences in the conductivity and permittivity values of human skin (on which the FEM model was based – solid line) compared to pig skin (on which validation measurements were made – shaded area representing ± 1 SD of n=8 measurements). Pig skin at the abdomen lacks hair follicles and therefore sweat glands. This lowers the conductivity when compared to human skin, which has sweat glands even in regions without hair follicles (Avci et al., 2013), and explains the lower DC components of the waveforms measured in pig cadavers compared to simulation. Skin permittivity is highly dependent on the outermost stratum corneum layer. The pig skin had a higher permittivity than the FEM model, and therefore more distinct capacitive components to the waveform. Supplementary Material 4 shows the same waveforms when the model skin conductivity was adjusted to be lower and permittivity values were adjusted to be higher to reflect the properties of pig skin and a good fit was attained. Furthermore, while the model predicts a 10% efficiency (at DC portions of V_TES_) in current transfer from I_TES_ to I_Nerve_, the measurements show a value closer to 7%. This discrepancy in the capture ratio may be explained by highly conductive edema in the surgical pocket and interfacing of the collectors with subcutaneous fat (inseparable from skin), instead of lying flush with lower conductivity skin, as in the model.

### Mechanism of charge transfer to the nerve

Despite the exponential capacitive waveforms in Fig. 4 (b-c), the main mechanism of charge transfer to the nerve is ohmic. The TES-tissue interface is highly capacitive, but once charge enters tissue, ohmic charge transfer dominates. This trend was investigated by setting the skin conductivity to 0 S/m while leaving the permittivity unchanged in the transcutaneous coupling FEM model. A transient simulation was run, and charge transferred to the nerve was calculated as area under the rectified I_Nerve_ curve. In the Fig. 4 (c) waveform with the fastest rise time, 29% of the charge transferred to the nerve was maintained when the conductivity of skin was set to 0 S/m and the only way for charge to cross the skin layer was as displacement current. Data are shown in Supplementary Material 5.

### Validated transcutaneous coupling model to investigate Injectrode system parameters

The validated FEM model, Fig. 2 (c), using the original literature conductivity and permittivity values representing human skin, was then used to further explore how waveform and geometric parameters affected system performance. Given the slight discrepancies between the absolute values in the model and swine cadaver measurements, expected differences in a live chronic experiment where scarring and healing occurs, and expected differences between swine and humans, the modeling results should be interpreted in terms of trends instead of absolute values.

#### Effect of collector size on charge transfer efficiency

After validating the FEM model with cadaver measurements, the model was used to further explore the parameter space of the Injectrode system on coupling efficiency. One key parameter explored was the ratio of the TES patch size to the collector size. Collector diameter was varied between 0.5 and 7 cm while holding the distance between the centers of the two collectors constant. Efficiency, or capture ratio (Gan and Prochazka, 2010), defined as I_Nerve_/I_TES_, was plotted in Fig 5. (a). The modeling suggests that efficiency increases with collector size until the collector diameter approaches the TES side length, after which efficiency decreases. This increase in efficiency as a function of collector diameter is expected as voltage directly under the TES electrode is roughly constant and increased collector area translates to a lower impedance interface with tissue and more current captured by the collector. However, when collector diameter exceeds TES electrode side length, it enters an area of tissue where the electric potential quickly drops off. The collector is a metallic conductor and hence forms an equipotential surface, where the collector voltage is defined by the lowest potential the collector contacts. Therefore, as collector diameter exceeds TES side length, the collector shunts current from under the TES electrodes to the edges of the collector where the electric potential of tissue is lower, and efficiency decreases. The trend in coupling efficiency with collector diameter was also demonstrated in the monopolar configuration. Collector size can be increased approximately up to the size of the surface electrode to increase current delivered to the deep target nerve.

#### Effect of high-frequency waveform on charge transfer efficiency

High-frequency waveforms are sometimes used in non-invasive neurostimulation devices with the hypothesis that they improve penetration depth of the delivered current (Nonis et al., 2017; Medina and Grill, 2014). Our model shows that charge transfer efficiency is lower when using a high-frequency (10 kHz) stimulation waveform when compared to a DC stimulation waveform (orange and blue traces, respectively, in Fig. 5). Tissue impedance appears lower at 10 kHz due to the addition of capacitive charge transfer, which makes the path to the deep target nerve through the Injectrode collectors becomes less preferential for current compared to travelling in tissue – lowering coupling efficiency.

**Figure 5:**
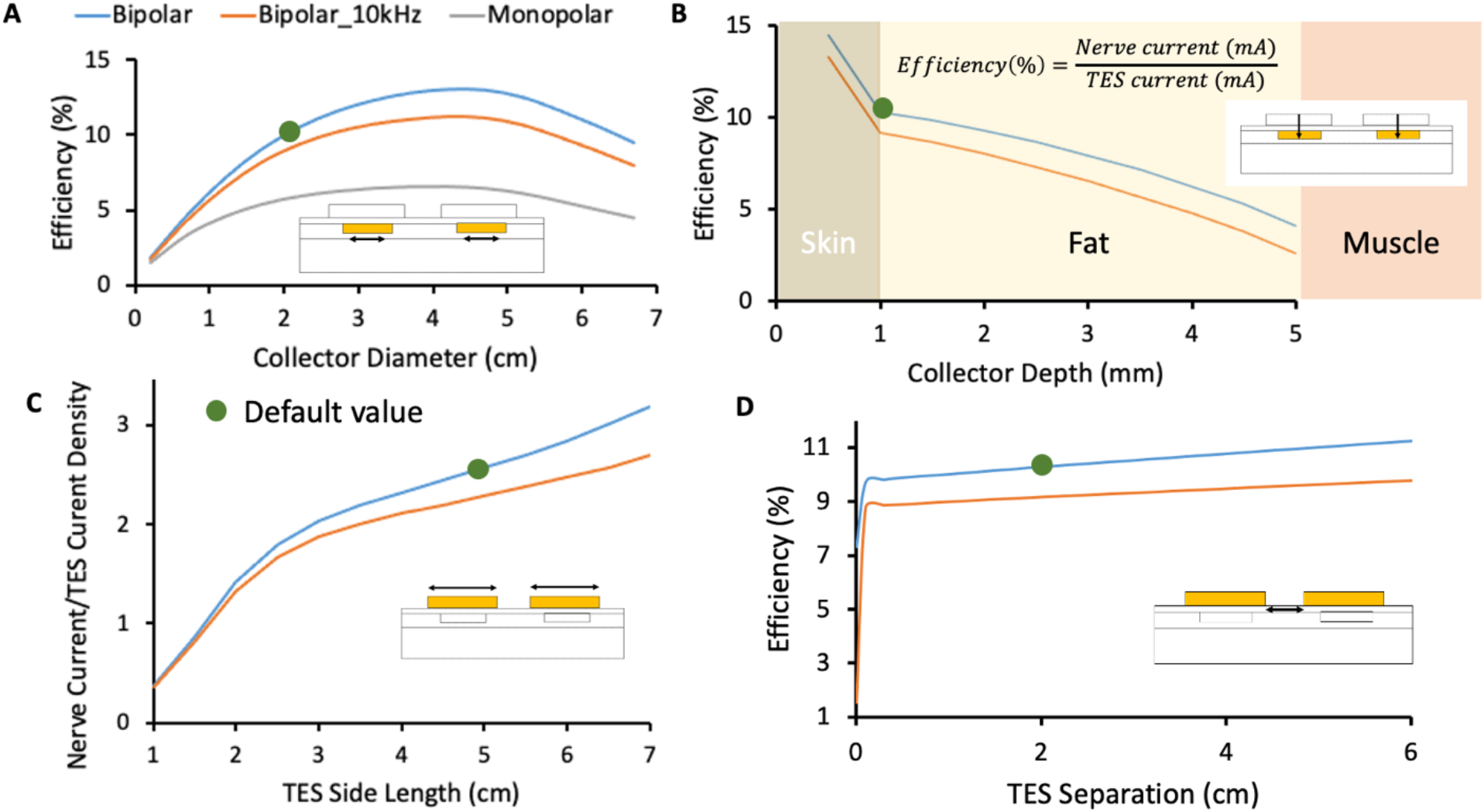
In this figure, blue and orange traces represent the Injectrode system in bipolar configuration with stimulation at DC and 10 kHz, respectively. The grey traces represent the Injectrode system in monopolar configuration with stimulation at DC. Green dots denote the default parameters used in the FEM model. **(a)** Change in efficiency with collector diameter. Optimal efficiency was achieved when the collector diameter approximately matched the surface electrode length. Tissue impedance is lower at higher frequency (orange trace at 10 kHz), which caused current to spread more and decreased capture efficiency. The current was more volumetrically contained with a bipolar setup (blue trace compared to monopolar grey trace). **(b)** Efficiency was highest closest to the surface electrodes and dropped quickest in the least conductive skin layer. **(c)** The ratio of I_Nerve_ to TES current density increased for larger surface electrode sizes. I_Nerve_ is a proxy for on-target recruitment of the deep nerve and surface current density is a proxy for recruitment of cutaneous off-target neural fibers. **(d)** Increasing separation between bipolar surface electrodes increased efficiency marginally by increasing the impedance of the leakage path from collector to collector compared to the low-impedance conduit formed by the Injectrode to the nerve. At small separations (<0.1 cm in this idealized model of dry skin) between the TES electrodes, current shunts superficially between the two electrodes and is not delivered deeper into tissue.

#### Effect of collector depth on charge transfer efficiency

A second key parameter that could impact the efficiency of charge transfer to the deep target nerve is the depth of the collector under the skin. To understand the sensitivity of transcutaneous charge transfer efficiency to collector depth, collector depth was varied from 0.5 to 5 mm – corresponding to the center of the 1 mm thick skin layer to the fat-muscle layer boundary. Fig. 5 (b) shows efficiency decreased with increased collector depth – most rapidly in the skin layer (14.5% to 10.3% over 0.5 mm) and then more gradually in the fat layer (10.3% to 9.9% over the first 0.5 mm). The decrease in efficiency was more rapid in the skin layer because skin is two orders of magnitude less conductive than fat and the electric potential drops off quickly in the skin layer with distance from the surface electrodes. The current through the 1 kΩ resistor connected between the two collectors, representing the deep nerve, is proportional to the voltage difference between the two collectors. As distance from the surface electrode increases, the voltage difference between the two collectors decreases. These data would suggest the collectors must be placed at the shallowest depth for highest efficiency. However, device extrusion (Zakhar et al., 2020; Uppal et al., 2021) and the position of sensory receptors in the skin (Crawford and Caterina, 2020) must also be considered to ensure that the collectors do not cause excessive pain.

#### Effect of surface electrode size on charge transfer efficiency

A critical parameter potentially impacting the ratio of on-versus off-target neural recruitment is the size of the TES electrodes. To investigate the effects of TES electrode size, edge-to-edge distance between the two surface electrodes was kept constant at 2 cm while the side length of the square surface electrodes was varied from 1 to 7 cm in the FEM model. The ratio of nerve current to surface electrode current density was plotted as a proxy for on-target nerve recruitment to off-target cutaneous fiber recruitment (Slopsema et al., 2018) in Fig. 5 (c), with the hypothesis that lower surface current densities generated by larger TES patches would reduce off-target cutaneous activation. Increasing the square surface electrode side length was found to improve the ratio of deep nerve current to surface electrode current density by ~8x when the TES side length was increased from 1 cm to 7 cm, suggesting a more favorable on-target deep nerve recruitment to off-target cutaneous fiber recruitment ratio at larger TES electrode sizes. Two effects are at play that make larger surface electrode sizes more suitable for preferential recruitment of on-target fibers. Firstly, larger surface electrode sizes result in lower TES-skin interface charge density for the same charge injected into tissue. Secondly, both collectors were centered under their respective surface electrode and the collector-to-collector separation increased with increased surface electrode size. The impedance of the path current must travel through tissue between the two collectors increased and the alternative path provided through the collectors and 1 kΩ resistor became more favorable, with more current directed to the deep target nerve. The model suggests that larger surface electrode sizes results in increased deep target nerve activation while decreasing paresthesia or pain caused by recruitment of off-target cutaneous fibers.

#### Effect of bipolar surface electrode separation on charge transfer efficiency

Lastly, a parameter impacting efficiency of charge transfer from the surface electrodes to the deep target nerve, but not as sensitively as the previous three parameters presented, is the separation between the two TES electrodes. The edge-to-edge separation between the two surface electrodes was varied from 0 to 6 cm and results are shown in Fig. 5 (d). Both collectors were kept centered under their respective surface electrodes. A minimum separation of 0.1 cm (smallest separation investigated) is required to prevent the surface electrodes from ‘shorting’ and shunting current superficially. Under more realistic conditions, such as sweating and the presence of blood vessels in skin (Khadka and Bikson, 2020), where skin impedance is drastically lowered, it is possible that shunting of current superficially between the two surface electrodes will be prominent at separations much larger than 0.1 cm.

However, this was not investigated. After that minimum separation, increased TES separation led to slightly increased efficiency (10% at 1 cm separation to 11% at 5 cm separation) as the path for current to flow through tissue between the two collectors increased in impedance. The current preferentially routed through the comparatively lower impedance pathway of the collectors and 1 kΩ resistor and more current was directed through the deep target nerve. These data suggest that surface electrode separation should be increased for minor gains in efficiency. However, this increased electrode separation would require a longer wire to be tunneled from the deep target nerve, where the Injectrode interfaces with the nerve, to the subcutaneous collectors – increasing injection complexity and volume of injected material.

### Validated transcutaneous coupling model to investigate patient-dependent parameters

A critical limitation of current TES therapies is that they are applied at home by an untrained user, without consideration of local anatomy, which affects current flow and neural activation (Zander et al., 2020). A goal of the Injectrode system is to reduce the sensitivity of existing TES therapies to patch placement by an untrained user and other expected intra- and inter-subject differences. Furthermore, an implanted neurostimulation device typically assures extent of neural recruitment by precisely controlling the current or voltage delivered to the nerve at the electrode-nerve interface. In the case of the Injectrode system, power delivered at the surface electrode-skin interface is regulated while the energy delivered at the Injectrode-nerve interface is not as precisely regulated. With increased transcutaneous coupling efficiency, the same surface electrode current would result in more current at the deep on-target nerve. The subject’s ability to ‘feel’ increased activation of the deep nerve via superficial sensation is minimized. Increased recruitment of an on-target autonomic nerve, for example, may not result in a sensation noticeable to the subject, yet still affect a physiological function unbeknownst to the patient. It is therefore crucial to understand the effect of variations in relevant parameters on coupling efficiency from the surface electrode to the deep target nerve. The FEM model was used to investigate coupling efficiency of the Injectrode system to expected inter- and intra-subject variations: placement of surface electrode relative to injected collector, skin thickness, and tissue conductivities and permittivities. Both current- and voltage-controlled waveforms were investigated and plotted in Fig. 6.

**Figure 6:**
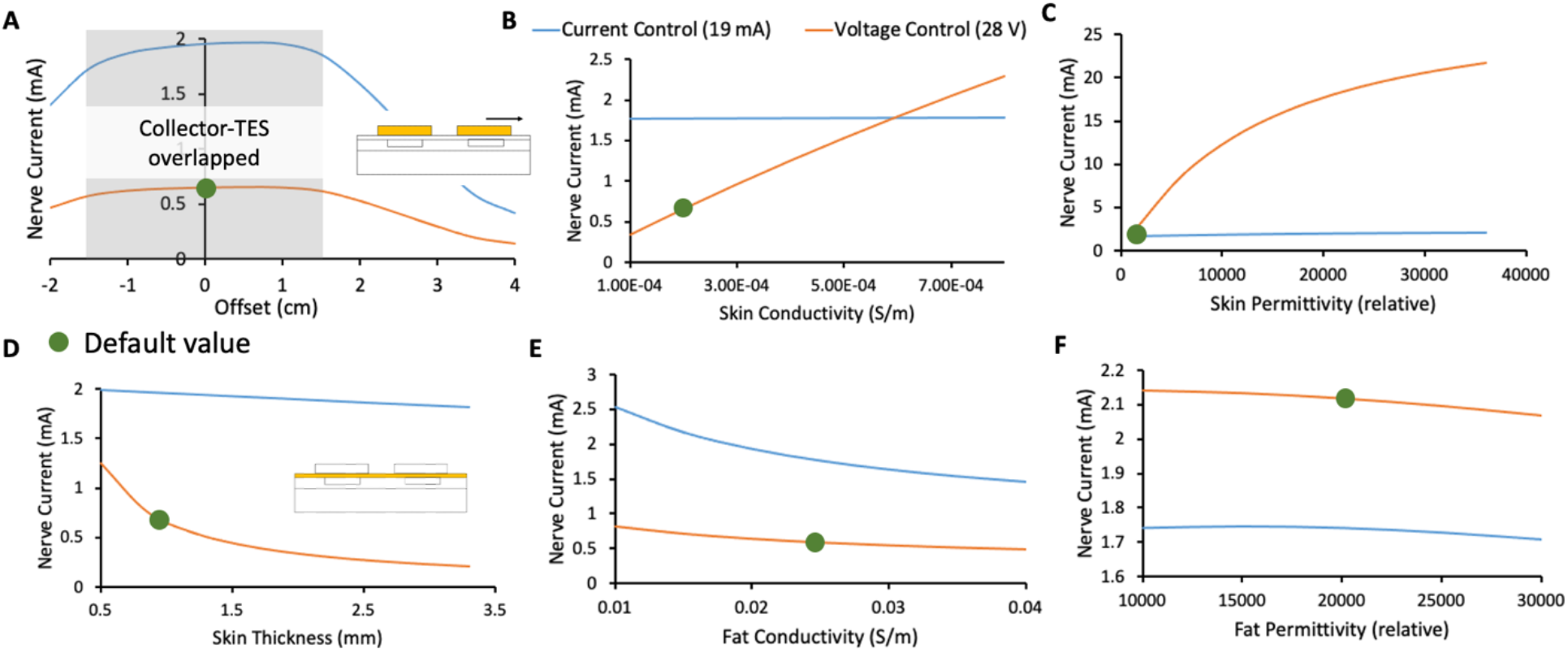
In this figure, orange and blue traces represent the deep target nerve current in response to voltage-controlled and current-controlled stimulation, respectively. Green dots denote the default parameters used in the FEM model. **(a)** Deep target nerve current is stable while the surface electrode completely overlaps the collector. **(b)** Deep target nerve current is more stable to variations in skin conductivity under current-controlled stimulation relative to voltage-controlled stimulation. **(c)** Deep target nerve current is more stable to variations in skin permittivity under current-controlled stimulation relative to voltage-controlled stimulation. **(d)** Deep target nerve current is more stable to variations in skin thickness under current-controlled stimulation relative to voltage-controlled stimulation. **(e)** Deep target nerve current is somewhat more stable to variations in fat conductivity under voltage-controlled stimulation relative to current-controlled stimulation. **(f)** Deep target nerve current is not sensitive to variations in fat permittivity.

#### Effect of collector and surface electrode offset on charge transfer efficiency

Fig. 6 (a) shows the stability in nerve current when a surface electrode is offset relative to its collector. Shaded in grey is the area where the collector remains entirely under the surface electrode, and the nerve current is stable. Nerve current drops rapidly when the surface electrode no longer overlaps the collector completely. Larger surface electrodes would be more tolerant to expected variations in reapplication, especially when done at home, along with the previously observed advantage in preferential on-target neural activation. Permanent markings on the skin or an automated electro impedance tomography (EIT) based system (Ansory et al., 2018) could also be used to guide at-home placement of the surface electrodes.

#### Effect of skin thickness and tissue electrical properties on charge transfer efficiency

In Fig 6. (b-d), the FEM model shows that current-controlled stimulation is stable to variations in skin thickness, conductivity, and permittivity, while voltage-controlled stimulation is highly susceptible to these variations. Skin thickness is expected to vary widely based on body location (Sandy-Møller et al., 2003) and age (Hoffmann et al., 1994; Neerken et al., 2004). Skin conductivity is also expected to vary widely based on sweating, weather, and skin preparation (Tronstad et al., 2010). Particularly, skin permittivity is based largely on the outermost thin stratum corneum layer and is expected to vary based on skin preparation (Tronstad et al., 2010). Skin preparation should be standardized and noted during clinical trials and replicated appropriately in clinical practice. On the other hand, nerve current during voltage-controlled stimulation was slightly more stable with variation in fat conductivity (Fig. 6 (e)). However, due to the reasons noted, it is expected that inter- and intra-subject variations in skin will be greater than fat. Given the critical safety and efficacy concerns surrounding a stable stimulation current delivered at the nerve (discussed earlier), and the larger expected variation in skin properties compared to fat properties, a current-controlled surface electrode stimulation may be preferred.

The analysis presented in this section creates an understanding of variations in the deep target nerve stimulation current with expected changes in patient-dependent parameters (e.g., surface electrode placement, tissue thickness, skin preparation, tissue electrical properties) during use of the Injectrode system. These analyses create a foundation for the Injectrode system to be designed to be more stable to expected patient-dependent changes, and thus improve device safety and efficacy for a wider patient population.

### Effect of the Injectrode on vagal axon activation (biophysical model)

The goal of the Injectrode is to facilitate the activation of deep neural targets using TES. To investigate if the Injectrode system can achieve this goal, we examined how including the Injectrode system components (i.e., the subcutaneous collector, Injectrode lead, and Injectrode nerve interface) affected the activation thresholds of target Aβ-axons in the vagus nerve compared to TES alone. Including the Injectrode system components lowered on-target vagus Aβ-axon activation thresholds by one to two orders of magnitude, regardless of patch size (Fig. 7 (b)). When using a 2×2 cm TES patch with the Injectrode system, the median vagus Aβ-axon activation threshold was 2.76 mA, compared to 57.69 mA with TES alone. This estimate of ~57 mA of stimulation current to activate Aβ-axons in the human cervical vagus is in line with a recent *in vivo* study showing ~34 mA was required to non-invasively activate A-fibers in a rat, where the cervical vagus is at a more superficial depth (Bucksot et al., 2020). When using a 3×3 cm TES patch with the Injectrode system, the median vagus Aβ-axon activation threshold was 4.12 mA, compared to 72.53 mA with TES alone. When using a 5×5 cm TES patch, the median vagus Aβ-axon activation threshold was 8.58 mA when using the Injectrode system, compared to 117.38 mA with TES alone. Vagus Aβ-axon thresholds were always lower when using the Injectrode system compared to TES alone.

**Figure 7:**
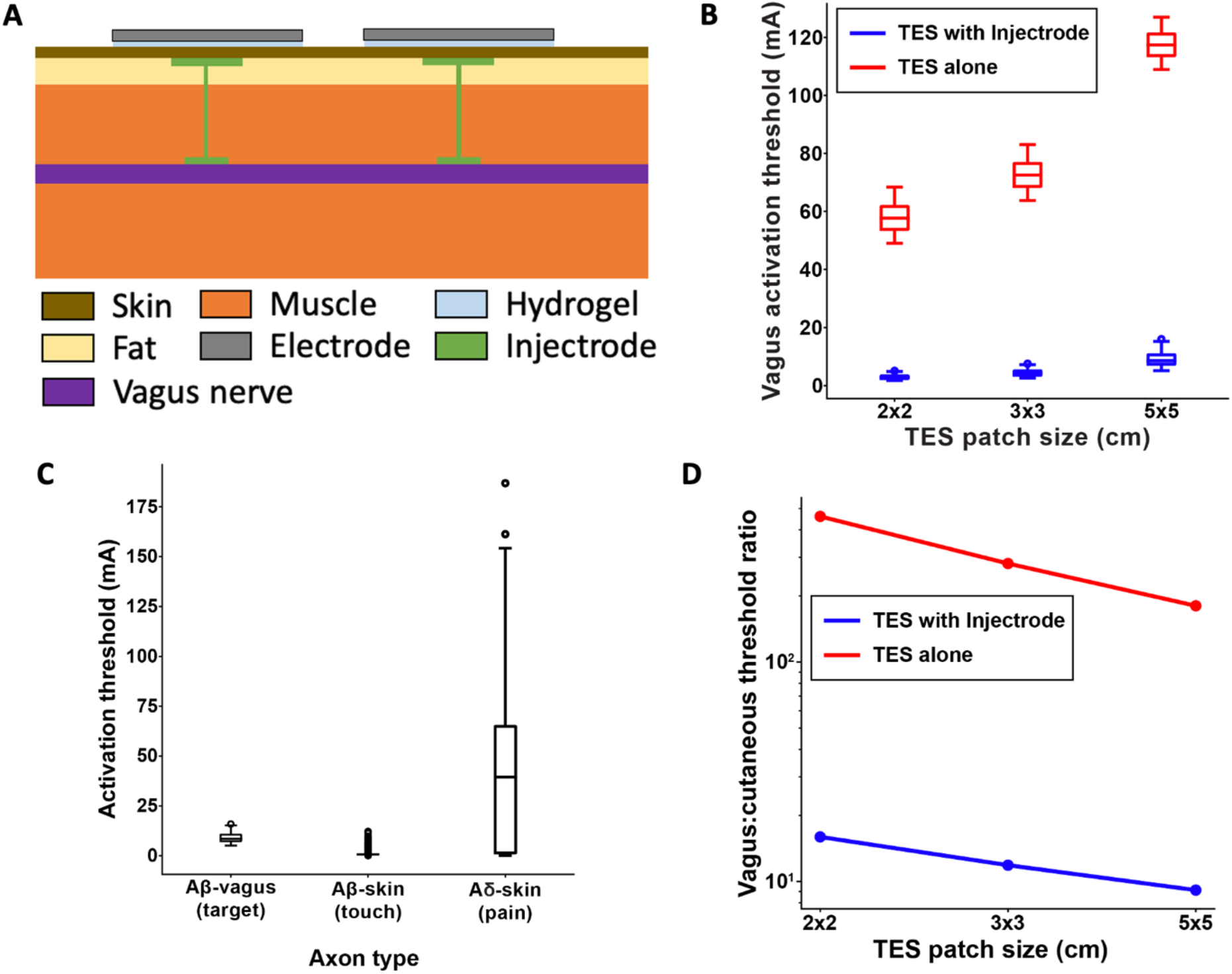
**(a)** Full FEM model used in the biophysical study. The 1 kΩ resistor between the two collectors in the simplified transcutaneous coupling FEM model was replaced with the vagus nerve, Injectrode connections down to the vagus nerve, and Injectrode interfaces with the vagus nerve. **(b)** Box plot showing that the Injectrode system reduced the current required to activate A*β* vagal fibers by more than an order of magnitude compared to using only surface electrodes. This large difference was seen across TES patches of different sizes. **(c)** Box plot of activation thresholds of on-target A*β* vagal fibers compared to off-target A*β* cutanoues fibers responsible for paresthesia and off-target A*δ* cutaneous fibers responsible for noxious sensations. **(d)** Investigating the effect of TES patch side length on cutaneous and vagus A*β*-fiber activation. Increasing the TES patch side length increased median thresholds, and this effect was less pronounced in vagal A*β* fibers, improving the ratio of A*β* vagal activation to A*β* cutaneous activation. The Injectrode system achieved preferential on-target activation one to two orders of magnitude better than using surface stimulation electrodes alone.

### Effect of the Injectrode system on on- and off-target neural recruitment (biophysical model)

It is possible that the Injectrode system could produce undesired activation of cutaneous afferents when using stimulation amplitudes necessary to activate target axons in deeper nerve structures. Therefore, we compared the activation thresholds of target Aβ-axons in the vagus nerve to the activation thresholds of off-target Aβ- and Aδ-cutaneous axons when using a 5×5 cm TES patch (Fig. 7 (c)). The median vagus Aβ-axon activation threshold was 8.58 mA. The median cutaneous Aβ-axon activation threshold was 0.73 mA. The median cutaneous Aδ-axon activation threshold was 39.41 mA. Generally, the distributions of target vagus Aβ-axons thresholds and cutaneous Aβ-axon thresholds overlapped more than the distributions of target vagus Aβ-axons thresholds and cutaneous Aδ-axon thresholds when using the Injectrode system. Using only 5×5 cm TES patches (without the Injectrode system), the median vagus Aβ-axon activation threshold was 117.38 mA – much higher than the median activation threshold of cutaneous Aβ- and Aδ-axons.

### Effect of surface electrode size on relative activation of on-target vagal and off-target cutaneous axons (biophysical model)

Results from the simplified transcutaneous coupling FEM model showed that increased TES patch sizes increased the ratio of current in the deep target nerve to current density at the surface electrodes (Fig. 5 (c)), which suggests an improved ratio of on-target neural activation in the deep nerve to off-target cutaneous activation. We used the field-cable model to verify whether increasing TES patch size increases the ratio of neural activation in the vagus Aβ-axons compared to cutaneous Aβ-axons. We were particularly interested in how changing TES patch size may lead to paresthetic percepts resulting from TES while achieving neural activation in the vagus nerve. Therefore, we examined how increasing TES patch size affected the ratio of the median activation thresholds of the ten most excitable vagus axons and the ten most excitable cutaneous axons (i.e., the axons with the lowest activation thresholds). Though we are only presenting threshold ratios for the ten axons with the lowest thresholds, this trend held for the same analysis when considering more than 50 of the most excitable axons (data not shown).

When using a TES electrode side length of 2 cm, the ratio of vagus-to-cutaneous median thresholds was 15.97 when using an Injectrode, compared to 460.66 when using TES alone. Increasing the TES electrode side length to 3 cm lowered the vagus-to-cutaneous median threshold ratio to 11.87 when using an Injectrode, compared to 281.03 when using TES alone. Further increasing the TES side length to 5 cm produced vagus-to-cutaneous threshold ratios of 9.14 and 180.37 when using an Injectrode and TES alone respectively. Overall, increasing TES side length reduced the median threshold ratio of axons in the vagus nerve compared to cutaneous afferents, suggesting that larger TES side lengths improve the engagement of on-target vagal axons while reducing off-target cutaneous axon activation. These results support the findings of the simplified transcutaneous coupling FEM model investigation. Additionally, the Injectrode reduced threshold ratios by greater than an order of magnitude compared to using only surface electrodes, further underscoring the utility of the Injectrode system in activating deep nerves during TES.

## Discussion

The goal of this study was to set up and use a validated FEM model to understand the transcutaneous energy delivery from surface electrodes to Injectrode collectors using low-frequency electric fields. The model was used to study waveform design and system parameters as well as sensitivity of the Injectrode system to expected inter- and intra-subject variations. Regarding expected variability in the deployment of the Injectrode and at-home application of the external TES electrodes, we found:

1. <u>Collector depth</u> to be a sensitive parameter affecting transcutaneous charge coupling efficiency
2. Optimal <u>collector size</u> to be approximately equal to the <u>external TES electrode size</u> and a sensitive parameter affecting transcutaneous charge coupling efficiency
3. Increasing external TES electrode size to improve preferential recruitment of on-target deep neurons
4. <u>Bipolar TES separation</u> to be an insensitive parameter on transcutaneous charge coupling efficiency, while greater than a minimum separation of 0.1 cm (in idealized model of dry skin)
5. <u>Placement of external TES electrodes</u> to be an insensitive parameter on transcutaneous charge coupling efficiency, while the Injectrode collector is completely overlapped by the TES electrode
6. <u>Skin thickness, conductivity, and permittivity</u> to be insensitive parameters on transcutaneous charge coupling efficiency under current-controlled stimulation, but sensitive under voltage-controlled stimulation

### High-frequency carrier waveform

High-frequency waveforms have been proposed to transfer additional charge to the nerve while keeping the absolute amplitude of voltage applied across the surface electrodes constant. A sinusoidal 10 kHz carrier waveform was considered here as a method to get more current from surface electrodes to deep target nerves. Previous works showed that a high-frequency voltage-controlled waveform resulted in more current to deeper neural structures (Medina and Grill, 2014; Bucksot et al., 2020), although these studies concluded that the biophysics of neural activation would not necessarily lead to increased neural recruitment due to a filtering effect of the neural membrane on the high-frequency waveform (Medina and Grill, 2014; Medina and Grill, 2016). Results with the Injectrode system in Fig. 5 suggest that a high-frequency carrier would lead to less efficient coupling. For the same current delivered at the surface electrodes, less current was captured by the collectors and routed to the deep target nerve. This trend occurs because the effective tissue impedance decreases at high frequencies and provides a lower-impedance path for current to travel between the two collectors as opposed to the intended Injectrode route from the collector through the deep target nerve.

In both the TES only and TES with Injectrode situation, a higher frequency waveform would deliver additional current at the same voltage (Medina and Grill, 2014); however, the increased current would mean that off-target activation of cutaneous fibers would also increase (Fig. 5 (c)). This finding was also observed in Bucksot et al. (2020), where they showed that high-frequency sinusoidal waveforms required larger voltage amplitudes to non-invasively stimulate the vagus nerve in rats and that the ratio of on-target to off-target neural recruitment did not change with frequency (Bucksot et al., 2020). It is possible that high-frequency waveforms may act to suppress neural activation close to the electrode akin to a high-frequency nerve block (Mirzakhalili et al., 2020); however, this is speculative and evidence to the contrary exists (Medina and Grill 2014; Mirzakhalili et al., 2020).

### Waveform design for the Injectrode system

The waveform used in the Injectrode system may be optimized to increase on-target neural recruitment and minimize off-target neural activation, knowing the dual mechanisms of ohmic and capacitive charge transfer, utility of high-frequency carrier waveforms, underlying tissue composition, and expected inter- and intra-subject variabilities. Given the large dependence of nerve current on skin thickness, conductivity, and permittivity, a current-controlled waveform should be selected. This trend occurs because a current-controlled waveform delivers more energy when resistance is high and less energy when resistance is low, keeping the spread of energy more consistent despite variations in tissue impedance. If larger variations in the subdermal fat conductivity are expected, then a voltage-controlled waveform may be considered (Fig. 6 (e)). Overall, the current-controlled waveforms were found to more consistently regulate the current delivered to the nerve given expected variability in tissue thicknesses and electrical properties.

In general, a high-frequency carrier waveform is unlikely to be useful as it decreases coupling efficiency and may not improve the ratio of on-target to off-target neural fiber activation (Fig. 5 (c)) (Bucksot et al., 2020). In some situations, where skin resistance is extremely high and ohmic charge transfer across the skin is challenging, high-frequency waveform components may still be useful to transfer charge by displacement current instead. A voltage-controlled waveform delivering pulses with controlled rise times, as in Fig. 4 (b-c), could be used for this purpose. The ability to deliver additional current using displacement current (Fig. 4 (b-c)) should also be kept in mind when selecting therapeutic targets for the Injectrode system. For example, the high-frequency sinusoidal waveforms characteristic of certain nerve blocks would lend naturally to the high dV_TES_/dt that generates displacement currents.

### Geometry design of Injectrode system

Similarly, the geometry of the Injectrode systems, such as the size of the collector and surface electrodes and collector to collector separation, may be optimized to achieve on-target deep fiber activation while minimizing off-target superficial fiber activation. Maximizing surface electrode size was shown to improve on-target vs. off-target neural fiber recruitment (Fig. 5 (c) and Fig. 7 (d)). Increased surface electrode size also increased the tolerance of the nerve current to differences in placement of the surface electrode relative to the collector (Fig. 6 (a)).

Our results form a conceptual basis for and complement earlier experimental work by Gan and Prochazka (Gan and Prochazka, 2007; Gan and Prochazka, 2010). They implemented TES surface electrodes, subcutaneous stainless-steel discs as collectors, and insulated wires running to a nerve cuff interfacing at the common peroneal nerve in rabbits to investigate parameters such as capture ratio (i.e., efficiency) and threshold current for a motor response from the animals. Their experimental trends match the theoretical findings presented here. They report higher capture ratios of up to ~0.4 (i.e., 40% efficiency). The higher efficiency of their system may be due to the lower impedance of the conductive path to the nerve and the tissue geometry of the rabbit compared to the pig.

In clinical deployment of the Injectrode, ultrasound and electrical impedance tomography (EIT) data may be used to develop subject-specific models of activation. The therapy could be personalized using ultrasound data gathered in the doctor’s office – which may already be used to guide the injection of Injectrode at the on-target nerve – and extracting tissue thickness data to tailor the geometry and waveform parameters. EIT measurements may also be collected to select an appropriate waveform. Continuous EIT monitoring may be done using the surface electrodes to understand the electrical properties of the underlying tissue and Injectrode as they change on the chronic time scale (Cooper et al., 2011) as well as with body position (Kim et al., 2013). Motion artifact on the EIT recordings from stimulation-evoked muscle activation may also provide data to the presence of on- and off-target effects.

In this study, we focused our investigation on the transcutaneous charge coupling efficiency and did not investigate parameters of the Injectrode-nerve interface, such as ‘cuff’ length and nerve interface position relative to TES surface patches, which affect local neural fiber excitation (Roointan et al., 2020). Recent work investigated the localized ‘virtual bipole’ created when a metal cuff is in contact with neural tissue (Roointan et al., 2020). The work by Roointan and colleagues suggests that a longer length of Injectrode on the nerve, higher Injectrode conductivity, and a slight offset of the Injectrode-nerve interface from the center of the surface stimulation electrodes will minimize the current threshold required to activate on-target neural fibers.

### Selection of tissue electrical properties for FEM model and their limitations

Measuring the electrical properties of tissue is challenging, and the results are sensitive to the measurement methods and tissue preparation. Consequently, widely varying values of tissue conductivity and permittivity are reported in the literature. Measurements at low frequencies (used in this study) are even more challenging. The values in this low-frequency range, taken from the Gabriel et al. (1996b) database, are inaccurate up to 25%, as quoted by the authors (Gabriel et al., 1996a), due to the two-electrode method used to make the measurements. The two-electrode method at low frequencies results in a high impedance across the electrode-tissue interface and the addition of a substantial electrochemical potential drop to the recording. The study attempted to compensate with calibrations in saline solutions. In general, the values of fat and muscle conductivity are more reliable and consistent across studies compared to skin conductivity and permittivity values, which vary widely between studies.

Large variances in published values of skin electrical properties are due to the skin being composed of several layers. The outer most is the ~30 *μ*m thin stratum corneum (Mørch et al., 2011), a layer of dead skin with low conductivity and high permittivity (Tronstad et al., 2010). Depending on tissue preparation and handling, this outer layer can become fractured or peel off. Furthermore, the frequency of the waveform used for measurements dictates the effective depth at which the measurement is being taken (Tronstad et al., 2010). This issue is especially pertinent at low frequencies, where the effective depth is shallow, and sampling is prominent in the outermost stratum corneum layer (Tronstad et al., 2010).

Furthermore, the electrical properties of skin change under the application of surface electrodes (Tronstad et al., 2010; Vargas Luna et al., 2015) and the delivery of electrical stimulation (Chizmadzhev et al., 1998). Gel used in the application of surface electrodes enters pores in the skin, lowering skin impedance (Tronstad et al., 2010; Vargas Luna et al., 2015). A similar effect is observed due to sweating under the surface electrodes after prolonged use (Tronstad et al., 2010; Vargas Luna et al., 2015). The delivery of an electric current during TES sets up an electric field across the skin, possibly resulting in electroporation of the skin (Chizmadzhev et al., 1998). Application of >30 V across the skin causes breakdown of the skin layers and creation of additional pathways of conduction through the skin, resulting in lower impedance (Chizmadzhev et al., 1998). Electroporation lasts for minutes to hours and may be irreversible if too high of a voltage is applied (Chizmadzhev et al., 1998). Skin impedance is also a function of the voltage applied even before electroporation occurs (Chizmadzhev et al., 1998).

### Effects of hydrogel

Hydrogel applied at the skin-TES electrode boundary plays an important role in limiting the coupling current from the surface electrodes. The resistive hydrogel layer increases the RC time constant of charging and decreases current spikes in I_TES_ (data not shown). Furthermore, the hydrogel distributes the current density across the surface electrode-skin interface and ensures it is not concentrated along the edges of the surface electrodes (Khadka and Bikson, 2020). High current density at the edges would result in earlier cutaneous fiber activation. Conductivity of hydrogels as well as other surface electrode material, including more reusable options, should be further studied to optimize the Injectrode system.

### Implications to TES modeling

Our FEM analysis suggests that it may be important to consider tissue permittivity in standard TES modeling – even without the Injectrode collectors. Tissue permittivity is particularly important when high-frequency TES voltage components are present, such as in voltage-controlled pulses and high-frequency waveforms (e.g., in the gammaCore device (Nonis et al., 2017)). Charge relaxation time is often used to justify that the bulk RC time constants of tissues under consideration are too short for displacement current to be considered a significant factor (Zhu et al., 2017). However, sometimes charge relaxation times are only calculated for the nerve cell membrane, with the implication that the cell membrane will act as an RC filter to the neurons within, preventing high-frequency components from depolarizing the nerve and initiating action potentials. On the other hand, calculations of charge relaxation time for skin, fat, and muscle provide an imperative to consider permittivity (Gabriel et al., 1996b). Given that there are many types of sodium channels with different dynamics and that non-neural glial cells also modulate neural activity (Abdo et al., 2019), there is a possibility that charge due to displacement current may modulate neural function. Furthermore, high frequencies may not be filtered at the sensory receptors themselves, which are located at neuron endings and are more superficial and closer to the surface stimulation electrodes, experiencing higher potentials. The sensory receptor-to-neuron transitions and rapidly changing conductivity across skin layers also creates a unique set of boundaries for changing the activation function (i.e., second-order spatial derivative of the extracellular potentials) and thereby inducing neural activation. Therefore, permittivity may be important to consider for standard TES modeling, especially in high-frequency voltage-controlled waveforms.

Pertinently, TES modeling studies sometimes consider only the deep on-target nerve (Mourdoukoutas et al., 2018) and ignore the superficial and cutaneous neural structures that lie between the surface stimulation electrodes and the deep target nerve. Our results suggest that these more superficial off-target neural structures experience higher potentials from the stimulation electrodes and can have lower activation thresholds than the deep target nerve (Fig. 7 (c)). Recruitment of these superficial off-target structures can cause painful sides effects and limit the stimulation dose to sub-therapeutic levels. Our work supports the observation that off-target effects of the gammaCore device (visible as lip curl) are likely due to activation of the superficial cervical branch of the facial nerve, which runs in the neck under the stimulation electrodes and innervates the platysma muscle (Nonis et al., 2017).

Our results also suggest that the use of a voltage-controlled high-frequency carrier waveform, implemented in the gammaCore device (Nonis et al., 2017), does increase the current that reaches deeper tissue. However, it also increases overall current spread and decreases the portion of charge delivered to the on-target neural tissue (Fig. 5 (c)). Therefore, use of the high-frequency waveform may simultaneously increase off-target neural activation, as demonstrated in rat experiments (Bucksot et al., 2020).

### Implications of biophysical modeling of Injectrode system

We used a field-cable modeling approach to investigate how the Injectrode system affects the activation of on-target axons in deep nerve structures compared to off-target axons in the skin. First, we examined whether the presence of the Injectrode components lowered activation thresholds of on-target vagal neurons compared to TES alone. The median vagus axon activation thresholds were more than ten times lower when using the Injectrode system compared to TES without Injectrode components (Fig. 7 (b)). This trend suggests that the Injectrode system can dramatically reduce the activation threshold of axons in deep nerves and underlines the potential utility of the Injectrode as a minimally invasive clinical strategy to stimulate deep neural targets. However, there still are potential off-target effects of the Injectrode system, such as activation of nociceptive cutaneous afferents, which must be mitigated to ensure patient comfort.

Next, we investigated potential on- and off-target effects of the Injectrode system by comparing the activation thresholds of target Aβ-axons in the vagus nerve to the activation thresholds of Aβ- and Aδ-cutaneous axons (Fig. 7 (c)). Generally, Aβ-cutaneous axons had the lowest activation thresholds, likely because of their large diameter and proximity to the surface stimulation electrodes. However, there is noticeable overlap in the distributions of Aβ-cutaneous and vagus thresholds. The cutaneous Aδ-axon threshold distribution was more variable than the other two distributions, and the median Aδ-axon threshold was considerably higher than the Aβ-vagus and cutaneous median thresholds. These results suggest that the Injectrode system may produce innocuous paresthesias as a side effect while activating deep nerve targets, without producing painful cutaneous sensations mediated by Aδ-fibers. In contrast, TES without the aid of the Injectrode had a median Aβ-vagus threshold that was much higher than the median threshold of cutaneous Aβ- and Aδ-axons. This result suggests that attempts to activate Aβ-vagus fibers using only surface stimulation electrodes may result in widespread activation of painful cutaneous Aδ-axons.

Lastly, we analyzed how design parameters, such as the TES patch size, affected the relative activation of cutaneous and vagus axons via the Injectrode system (Fig. 7 (d)). In general, increasing TES patch size increased the activation thresholds of both cutaneous and vagus axons. However, the increase in cutaneous activation thresholds was larger than the increase in target vagus activation thresholds, and this effect became more pronounced as the patch side length increased. Therefore, using larger TES surface patches may reduce the off-target activation of cutaneous afferents relative to on-target activation of deep nerve structures.

### General limitations of study

While the FEM model used in this study represented the skin, fat, and muscle as distinct domains, they were seen to be fused together in the swine cadaver studies. In particular, the skin and subcutaneous fat layers were inseparable and a source of error when measuring the thickness of skin, which is also the distance between the surface electrode and collector. Sensitivity analysis presented in Fig. 6 (d) shows that the current delivered to the nerve under voltage-controlled stimulation is sensitive to skin thickness. Ultrasound imaging may be used in future studies to more accurately quantify skin and fat thicknesses.

As explained in the Methods section, a 2.1 cm diameter stainless-steel disc was used as a consistent representation of the Injectrode collector. While the stainless-steel showed similar performance in the acute cadaver experimentation, it is possible that the porosity of the Injectrode will allow revascularization during chronic use, changing its electrical properties and corresponding performance compared to a stainless-steel disc. Chronic performance of the transcutaneous charge coupling mechanisms needs to be further investigated.

We did not consider off-target neural recruitment by electric current from the Injectrode leads connecting the Injectrode collectors to the nerve interface (Fig. 7 (a)). In the implementation of the Injectrode where the lead wires are insulated, this source of off-target neural recruitment is unlikely. However, if uninsulated leads are used to connect the collector to the neural interface, off-target recruitment of neurons by the leads must be considered.

A simplified FEM model was used in the transcutaneous charge coupling investigation to isolate the effects of system parameters on transcutaneous coupling between the surface electrodes and collectors. As part of the simplification, a 1 kΩ resistor was used to model the Injectrode path between the two collectors, based on EIS measurements from Trevathan et al. (2019). In reality, the impedance of the path between the two collectors varies as a function of frequency, voltage, and tissue properties. However, at the same time, the alternate conduction path through tissue (leakage through tissue between the two collectors) would also vary similarly and the general trends presented here are expected to hold.

An instrumentation limitation explains why the spikes in surface electrode current predicted by the FEM model are higher than the cadaver measurements. The Keithley DAQ 6510 used to collect surface electrode current measurements is bandwidth limited, recording −3 dB at 25 kHz (Keithley). This frequency response means that higher frequency components were attenuated in their measured amplitude. However, the voltage measurements, made using an oscilloscope, were not bandwidth limited for the frequency range under investigation.

We used a field-cable model to investigate how the Injectrode system activates on- and off-target axons. Though the FEM and multi-compartment axon models were both constructed using experimental data and previously validated models, there are several limitations to this approach. For example, we used a previously published, but simplified, morphology to represent cutaneous Aβ- and Aδ-axons (Tigerholm et al., 2019). The cutaneous Aβ-axon terminated in a passive node of Ranvier, while the cutaneous Aδ-axon terminated in a branching structure designed to mimic the nerve fiber density in human skin. The extent to which terminal branching morphologies affect the neural response to TES is currently unknown. Future studies should examine how the complexity of branching structures, as well as electrophysiological differences across different types of sensory terminals, affect the neural response to TES.

Finally, we distributed cutaneous afferent terminals beneath the TES patch on a grid that extended 5 mm beyond the edges of the TES patch. This implies that using larger TES edge lengths would sample from more cutaneous axons. Therefore, when comparing cutaneous activation across different TES patch sizes, we were not making a one-to-one comparison between modeling conditions and may be over- or under-estimating the extent of neural activation across patch sizes. For this reason, when examining the effect of patch size on cutaneous activation, we compared the median thresholds of the 10 cutaneous axons with the lowest thresholds to understand how increased patch size affects the most excitable axons. It is also unclear how many cutaneous axons need to be activated to produce a painful or non-painful percept. Some argue that multiple afferents must be activated to produce a perceptible sensation (Wall and McMahon, 1985), while others argue activation of a single axon can produce a percept (Torebjörk, 1987). Future modeling studies should be paired with psychophysical experiments to determine how many axons are needed to induce an innocuous or painful percept.

### Conclusion

The Injectrode system, a minimally invasive technology, may provide the capabilities to recruit deep on-target fibers while minimizing side effects from off-target neural activation. This modeling study provides a framework on which to optimize the design and deployment of the Injectrode system. We investigated the transcutaneous charge coupling efficiency and neural fiber selectivity of the Injectrode system using validated computational models. Our results suggest that the Injectrode system lowers the activation thresholds of deep on-target vagal fibers by more than an order of magnitude compared to using only surface stimulation electrodes. This reduction in activation thresholds makes it possible to activate vagal fibers with only innocuous recruitment of paresthesia inducing Aβ-cutaneous fibers. Meanwhile, surface stimulation applied without the Injectrode system, will likely recruit painful Aδ-cutaneous fibers before recruiting target deep vagal fibers. Exploration of the parameter space of the Injectrode system suggests that surface electrode and collector size can be selected to increase the current delivered to the deep target nerve while reducing surface electrode current density, a proxy for off-target cutaneous fiber activation. Voltage- and current-controlled waveforms were considered to assess how variability in anatomical parameters, such as tissue electrical properties and thicknesses, affects the performance of the Injectrode system. Current-controlled waveforms were found to be more stable to variations in the skin layer. High-frequency waveforms were also investigated but were unsuccessful in preferentially increasing on-target vs. off-target neural activation. Our work highlights the need to consider the activation of superficial off-target neural structures when targeting deep neural fibers with non-invasive electrical stimulation modalities.

## Acknowledgements

EJ for sourcing and delivering swine cadavers used in this study. Carly Frieders, Maria LaLuzerne, Rex Chin-Hao Chen, Ashlesha Deshmukh, and Danny Lam for reviewing drafts of this manuscript. Members of the Wisconsin Institute for Translational Neuroengineering (WITNe) for feedback during group meetings.

## References

Abdo, H., Calvo-Enrique, L., Lopez, J. M., Song, J., Zhang, M.-D., Usoskin, D., et al. (2019). Specialized cutaneous Schwann cells initiate pain sensation. Science 365, 695–699. doi:10.1126/science.aax6452.

Ansory, A., Prajitno, P., and Wijaya, S. K. (2018). Design and development of electrical impedance tomography system with 32 electrodes and microcontroller. AIP Conference Proceedings 1933, 040023. doi:10.1063/1.5023993.

Aristovich, K., Donega, M., Fjordbakk, C., Tarotin, I., Chapman, C. A. R., Viscasillas, J., et al. (2021). Modelbased geometrical optimisation and in vivo validation of a spatially selective multielectrode cuff array for vagus nerve neuromodulation. Journal of Neuroscience Methods 352, 109079. doi:10.1016/j.jneumeth.2021.109079.

Avci, P., Sadasivam, M., Gupta, A., De Melo, W. C., Huang, Y.-Y., Yin, R., et al. (2013). Animal models of skin disease for drug discovery. Expert Opinion on Drug Discovery 8, 331–355. doi:10.1517/17460441.2013.761202.

Bossetti, C. A., Birdno, M. J., and Grill, W. M. (2008). Analysis of the quasi-static approximation for calculating potentials generated by neural stimulation. J. Neural Eng. 5, 44–53. doi:10.1088/1741-2560/5/1/005.

Bucksot, J. E., Morales Castelan, K., Skipton, S. K., and Hays, S. A. (2020). Parametric characterization of the rat Hering-Breuer reflex evoked with implanted and non-invasive vagus nerve stimulation. Experimental Neurology 327, 113220. doi:10.1016/j.expneurol.2020.113220.

Butson, C. R., and McIntyre, C. C. (2005). Tissue and electrode capacitance reduce neural activation volumes during deep brain stimulation. Clinical Neurophysiology 116, 2490–2500. doi:10.1016/j.clinph.2005.06.023.

Carome, M. A. (2020). Implanted Spinal Cord Stimulators for Pain Relief. Public Citizen. doi: https://www.citizen.org/wp-content/uploads/2526200610Spinal-Cord-Stimulator-ReportFINAL.pdf?eType=EmailBlastContent&eId=765162c2-baeb-41db-a6ae-0694813ad96c

Chizmadzhev, Y. A., Indenbom, A. V., Kuzmin, P. I., Galichenko, S. V., Weaver, J. C., and Potts, R. O. (1998). Electrical Properties of Skin at Moderate Voltages: Contribution of Appendageal Macropores. Biophysical Journal 74, 14.

Cooper, G., Barker, A. T., Heller, B. W., Good, T., Kenney, L. P. J., and Howard, D. (2011). The use of hydrogel as an electrode–skin interface for electrode array FES applications. Medical Engineering & Physics 33, 967–972. doi:10.1016/j.medengphy.2011.03.008.

Crawford, L. K., and Caterina, M. J. (2020). Functional Anatomy of the Sensory Nervous System: Updates From the Neuroscience Bench. Toxicol Pathol 48, 174–189. doi:10.1177/0192623319869011.

Dalrymple, A. N., Ting, J. E., Bose, R., Trevathan, J. K., Nieuwoudt, S., Lempka, S. F., et al. (2021). Stimulation of the Dorsal Root Ganglion Using an Injectrode^®^. bioRxiv. doi:10.1101/2021.08.16.456553.

Keithley. DAQ6510 Data Acquisition and Logging, Multimeter System Datasheet. Rev 090617.

FDA. (1997). Design Control Guidance for Medical Device Manufacturers.

Foster, K. R., and Schwan, H. P. (1989). Dielectric Properties of Tissue and Biological Materials: A Critical Review. Critical Reviews in Biomedical Engineering. 17, 25–104.

Gabriel, S., Lau, R. W., and Gabriel, C. (1996a). The dielectric properties of biological tissues: II. Measurements in the frequency range 10 Hz to 20 GHz. Phys. Med. Biol. 41, 2251–2269. doi:10.1088/0031-9155/41/11/002.

Gabriel, S., Lau, R. W., and Gabriel, C. (1996b). The dielectric properties of biological tissues: III. Parametric models for the dielectric spectrum of tissues. Phys. Med. Biol. 41, 2271–2293. doi:10.1088/0031-9155/41/11/003.

Gan, L. S., Prochazka, A., Bornes, T. D., Denington, A. A., and Chan, K. M. (2007). A New Means of Transcutaneous Coupling for Neural Prostheses. IEEE Trans. Biomed. Eng. 54, 509–517. doi:10.1109/TBME.2006.886664.

Gan, L. S., and Prochazka, A. (2010). Properties of the Stimulus Router System, a Novel Neural Prosthesis. IEEE Trans. Biomed. Eng. 57, 450–459. doi:10.1109/TBME.2009.2031427.

Gaunt, R. A., and Prochazka, A. (2009). Transcutaneously Coupled, High-Frequency Electrical Stimulation of the Pudendal Nerve Blocks External Urethral Sphincter Contractions. Neurorehabil Neural Repair 23, 615–626. doi:10.1177/1545968308328723.

Graham, R. D., Bruns, T. M., Duan, B., and Lempka, S. F. (2019). Dorsal root ganglion stimulation for chronic pain modulates Aβ-fiber activity but not C-fiber activity: A computational modeling study. Clinical Neurophysiology 130, 941–951. doi:10.1016/j.clinph.2019.02.016.

Graham, R. D., Bruns, T. M., Duan, B., and Lempka, S. F. (2020). The Effect of Clinically Controllable Factors on Neural Activation During Dorsal Root Ganglion Stimulation. Neuromodulation: Technology at the Neural Interface, ner.13211. doi:10.1111/ner.13211.

He, F., Lycke, R., Ganji, M., Xie, C., and Luan, L. (2020). Ultraflexible Neural Electrodes for Long-Lasting Intracortical Recording. iScience 23, 101387. doi:10.1016/j.isci.2020.101387.

Hines, M. (2009). NEURON and Python. Front. Neuroinform. 3. doi:10.3389/neuro.11.001.2009.

Hines, M. L., and Carnevale, N. T. (1997). The NEURON Simulation Environment. Neural Computation 9, 1179–1209. doi:10.1162/neco.1997.9.6.1179.

Hoffmann, K., Stuucker, M., Dirschka, T., Goortz, S., El-Gammal, S., Dirting, K., et al. (1994). Twenty MHz B-scan sonography for visualization and skin thickness measurement of human skin. J Eur Acad Dermatol Venerol 3, 302–313. doi:10.1111/j.1468-3083.1994.tb00367.x.

Ilfeld, B. M., Plunkett, A., Vijjeswarapu, A. M., Hackworth, R., Dhanjal, S., Turan, A., et al. (2021). Percutaneous Peripheral Nerve Stimulation (Neuromodulation) for Postoperative Pain: A Randomized, Sham-controlled Pilot Study. Anesthesiology, 10.1097/ALN.0000000000003776. doi:10.1097/ALN.0000000000003776.

Khadka, N., and Bikson, M. (2020). Role of skin tissue layers and ultra-structure in transcutaneous electrical stimulation including tDCS. Phys. Med. Biol. doi:10.1088/1361-6560/abb7c1.

Kim, C. H. (2013). Importance of Axial Migration of Spinal CordStimulation Trial Leads with Position. Pain Phys 6; 16, E763–E768. doi:10.36076/ppj.2013/16/E763.

Krahl, S. (2012). Vagus nerve stimulation for epilepsy: A review of the peripheral mechanisms. Surg Neurol Int 3, 47. doi:10.4103/2152-7806.91610.

Kuhn, A., Keller, T., Lawrence, M., and Morari, M. (2009). A model for transcutaneous current stimulation: simulations and experiments. Med Biol Eng Comput 47, 279–289. doi:10.1007/s11517-008-0422-z.

Kumar, K., and Bishop, S. (2009). Financial impact of spinal cord stimulation on the healthcare budget: a comparative analysis of costs in Canada and the United States: Clinical article. SPI 10, 564–573. doi:10.3171/2009.2.SPINE0865.

Loeb, G. E. (2006). The BION devices: injectable interfaces with peripheral nerves and muscles. Neurosurg. Focus 20, 9.

Manson, G. A., Calvert, J. S., Ling, J., Tychhon, B., Ali, A., and Sayenko, D. G. (2020). The relationship between maximum tolerance and motor activation during transcutaneous spinal stimulation is unaffected by the carrier frequency or vibration. Physiol Rep 8. doi:10.14814/phy2.14397.

McIntyre, C. C., Richardson, A. G., and Grill, W. M. (2002). Modeling the Excitability of Mammalian Nerve Fibers: Influence of Afterpotentials on the Recovery Cycle. Journal of Neurophysiology 87, 995–1006. doi:10.1152/jn.00353.2001.

Medina, L. E., and Grill, W. M. (2014). Volume conductor model of transcutaneous electrical stimulation with kilohertz signals. J. Neural Eng. 11, 066012. doi:10.1088/1741-2560/11/6/066012.

Medina, L. E., and Grill, W. M. (2016). Nerve excitation using an amplitude-modulated signal with kilohertz-frequency carrier and non-zero offset. J NeuroEngineering Rehabil 13, 63. doi:10.1186/s12984-016-0171-4.

Mirzakhalili, E., Barra, B., Capogrosso, M., and Lempka, S. F. (2020). Biophysics of Temporal Interference Stimulation. Cell Systems 11, 557–572.e5. doi:10.1016/j.cels.2020.10.004.

Mørch, C. D., Hennings, K., and Andersen, O. K. (2011). Estimating nerve excitation thresholds to cutaneous electrical stimulation by finite element modeling combined with a stochastic branching nerve fiber model. Med Biol Eng Comput 49, 385–395. doi:10.1007/s11517-010-0725-8.

Mourdoukoutas, A. P., Truong, D. Q., Adair, D. K., Simon, B. J., and Bikson, M. (2018). High-Resolution Multi-Scale Computational Model for Non-Invasive Cervical Vagus Nerve Stimulation. Neuromodulation 21, 261–268. doi:10.1111/ner.12706.

Neerken, S., Lucassen, G. W., Bisschop, M. A., Lenderink, E., and Nuijs, T. (A. M.). (2004). Characterization of age-related effects in human skin: A comparative study that applies confocal laser scanning microscopy and optical coherence tomography. J. Biomed. Opt. 9, 274. doi:10.1117/1.1645795.

Nicolai, E. N., Settell, M. L., Knudsen, B. E., McConico, A. L., Gosink, B. A., Trevathan, J. K., et al. (2020). Sources of off-target effects of vagus nerve stimulation using the helical clinical lead in domestic pigs. J. Neural Eng. 17, 046017. doi:10.1088/1741-2552/ab9db8.

Nonis, R., D’Ostilio, K., Schoenen, J., and Magis, D. (2017). Evidence of activation of vagal afferents by non-invasive vagus nerve stimulation: An electrophysiological study in healthy volunteers. Cephalalgia 37, 1285–1293. doi:10.1177/0333102417717470.

Poulsen, A. H., Tigerholm, J., Meijs, S., Andersen, O. K., and Mørch, C. D. (2020). Comparison of existing electrode designs for preferential activation of cutaneous nociceptors. J. Neural Eng. doi:10.1088/1741-2552/ab85b1.

Roointan, S., Tovbis, D., Elder, C., and Yoo, P. B. (2020). Enhanced transcutaneous electrical nerve stimulation achieved by a localized virtual bipole: a computational study of human tibial nerve stimulation. J. Neural Eng. 17, 026041. doi:10.1088/1741-2552/ab85d3.

Sandby-Møller, J., Poulsen, T., and Wulf, H. C. (2003). Epidermal Thickness at Different Body Sites: Relationship to Age, Gender, Pigmentation, Blood Content, Skin Type and Smoking Habits. Acta Dermato-Venereologica 83, 410–413. doi:10.1080/00015550310015419.

Slopsema, J. P., Boss, J. M., Heyboer, L. A., Tobias, C. M., Draggoo, B. P., Finn, K. E., et al. (2018). Natural Sensations Evoked in Distal Extremities Using Surface Electrical Stimulation. TOBEJ 12, 1–15. doi:10.2174/1874120701812010001.

Störchle, P., Müller, W., Sengeis, M., Lackner, S., Holasek, S., and Fürhapter-Rieger, A. (2018). Measurement of mean subcutaneous fat thickness: eight standardised ultrasound sites compared to 216 randomly selected sites. Sci Rep 8, 16268. doi:10.1038/s41598-018-34213-0.

Tigerholm, J., Poulsen, A. H., Andersen, O. K., and Mørch, C. D. (2019). From Perception Threshold to Ion Channels—A Computational Study. Biophysical Journal 117, 281–295. doi:10.1016/j.bpj.2019.04.041.

Torebjörk, H. E., Vallbo, Å. B., and Ochoa, J. L. (1987). Intraneural Microstimulation in Man: Its Relation to Specificity of Tactile Sensations. Brain 110, 1509–1529. doi:10.1093/brain/110.6.1509.

Trevathan, J. K., Baumgart, I. W., Nicolai, E. N., Gosink, B. A., Asp, A. J., Settell, M. L., et al. (2019). An Injectable Neural Stimulation Electrode Made from an In-Body Curing Polymer/Metal Composite. Adv. Healthcare Mater. 8, 1900892. doi:10.1002/adhm.201900892.

Tronstad, C., Johnsen, G. K., Grimnes, S., and Martinsen, Ø. G. (2010). A study on electrode gels for skin conductance measurements. Physiol. Meas. 31, 1395–1410. doi:10.1088/0967-3334/31/10/008.

Uppal, P., Wright, T. B., Dahbour, L., Watterworth, B., Lee, S. J., Gattu, K., et al. Difficult removal of exposed peripheral nerve stimulator leads: a report of 2 cases. PR9 6, e946. doi:10.1097/PR9.0000000000000946.

Udo, E. O., Zuithoff, N. P. A., van Hemel, N. M., de Cock, C. C., Hendriks, T., Doevendans, P. A., et al. (2012). Incidence and predictors of short- and long-term complications in pacemaker therapy: The FOLLOWPACE study. Heart Rhythm 9, 728–735. doi:10.1016/j.hrthm.2011.12.014.

Vargas Luna, J. L., Krenn, M., Cortés Ramírez, J. A., and Mayr, W. (2015). Dynamic Impedance Model of the Skin-Electrode Interface for Transcutaneous Electrical Stimulation. PLoS ONE 10, e0125609. doi:10.1371/journal.pone.0125609.

Wall, P. D., and McMahon, S. B. (1985). Microneuronography and its Relation to Perceived Sensation. A Critical Review. Pain 21, 209–229. doi:10.1016/0304-3959(85)90086-7.

Wei, X. F., and Grill, W. M. (2009). Impedance characteristics of deep brain stimulation electrodes *in vitro* and *in vivo*. J. Neural Eng. 6, 046008. doi:10.1088/1741-2560/6/4/046008.

Zakhar, J. (2020). Un-LINQed: Spontaneous extrusion of newer generation implantable loop recorders. Indian Pacing and Electrophysiology Journal 20, 189–192. doi:10.1016/j.ipej.2020.04.005.

Zander, H. J., Graham, R. D., Anaya, C. J., and Lempka, S. F. (2020). Anatomical and technical factors affecting the neural response to epidural spinal cord stimulation. J. Neural Eng. 17, 036019. doi:10.1088/1741-2552/ab8fc4.

Zhu, K., Li, L., Wei, X., and Sui, X. (2017). A 3D Computational Model of Transcutaneous Electrical Nerve Stimulation for Estimating Aβ Tactile Nerve Fiber Excitability. Front. Neurosci. 11, 250. doi:10.3389/fnins.2017.00250.

