## Supplementary Material for "Augmented Transcutaneous Stimulation Using an Injectable Electrode"

**Supplementary Material 1 – Market overview of minimally invasive neuromodulation devices**

**Supplementary Material 2 – Electrochemical interfaces measurement and modeling**

**Supplementary Material 3 – Current-controlled waveform in swine cadaver**

**Supplementary Material 4 – Cadaver validation fitted with values representative of swine skin**

**Supplementary Material 5 – Mechanism of transcutaneous charge transfer**

**Supplementary Material 1 – Market overview of minimally invasive neuromodulation devices**

Table S1: Energy transfer methods used in minimally invasive neuromodulation devices

| **Fundamental technology** | **Company** |
|---|---|
| Thin percutaneous wire | SPR therapeutics |
| Ultrasound | Iota Biosciences (acquired) |
| Low frequency magnetic field (near-field, non-radiative) | SetPoint Medical |
| High frequency electromagnetic field (far/mid-field, radiative) | Neuspera, Nalu Medical, Stimwave |
| Low frequency electric field (non-radiative) | StimRouter, NeuronOff (this work) |

Examples of minimally invasive neuromodulation technologies cleared or approved by the FDA include the SPRINT PNS System, using percutaneous energy transfer on a thin wire, by SPR Therapeutics (Minneapolis, MN) for the treatment of acute pain. In addition, Freedom SCS System and Nalu Neurostimulation System, using radio frequency energy transfer, by Stimwave (Pampano Beach, FL) and Nalu Medical (Carlsbad, CA), respectively, for the treatment of chronic pain.

Several other minimally invasive therapies are in the development pipeline. SetPoint Medical (Valencia, CA) is developing a device, using near-field energy transfer, for inflammatory diseases such as Crohn's and Rheumatoid Arthritis. Neuspera (San Jose, CA) is working on a mid-field powered device for urinary urgency incontinence. Near-field, mid-field, and far-field energy transfer are classified based on the distance of energy transfer relative to the wavelength of the electromagnetic wave used. In near-field, the distance of energy transfer is short relative to the wavelength; electric and magnetic fields can exist independently of one another, and the waves are non-radiative. In far-field, the distance of energy transfer is long relative to the wavelength; electric and magnetic fields exist together, and the waves are radiative. Iota Biosciences (Berkeley, CA) is working on an ultrasound-powered neuromodulation technology platform.

**Supplementary Material 2 – Electrochemical interfaces measurement and modeling**

Electrochemical interfaces in the computational model were represented by surfaces that have both resistance and capacitance. The resistive and capacitive values used in the equivalent circuit were based on the empirically measured results described below.

<u>Hydrogel</u>

20 mA of current was sent through two hydrogel TES electrodes adhered to each other. Voltage required to deliver the 20 mA 300 $\mu$s pulse is shown in the red trace below. The faradaic component represents a resistance of ~50 Ohms. The non-faradaic portion is approximately 0.4 V in amplitude. These measurements were used to calculate the conductivity of hydrogel as 1.6 x 10^-2 S/m and relative permittivity of 1.4 x 10^6.

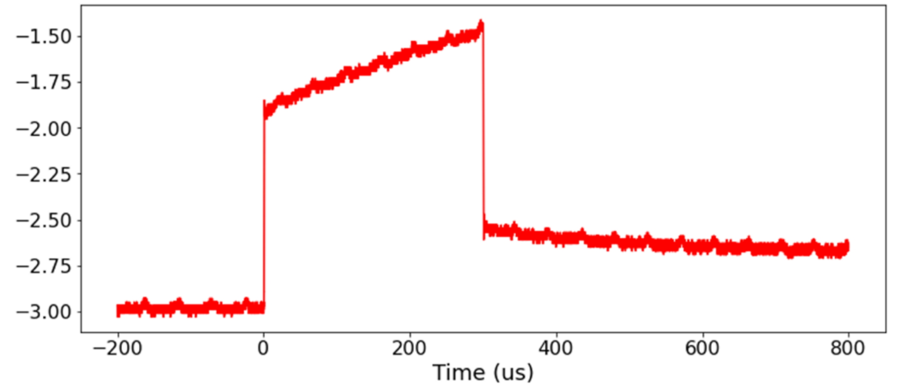


Figure S2a: Plot of voltage against time when a 20 mA current pulse was delivered to two TES hydrogel electrodes adhered to each other.

<u>Injectrode-tissue Interfaces</u>

Voltage was applied across two collectors immersed in saline solution and current drawn was measured. The applied voltage reflected what was expected at the collector based on preliminary cadaver measurements. Based on these measurements, the resistivity of the stainless-steel disc collector was calculated to be 6.9 x 10^-2 $\Omega.m^2$.

**Supplementary Material 3 – Current-controlled waveform in swine cadaver**

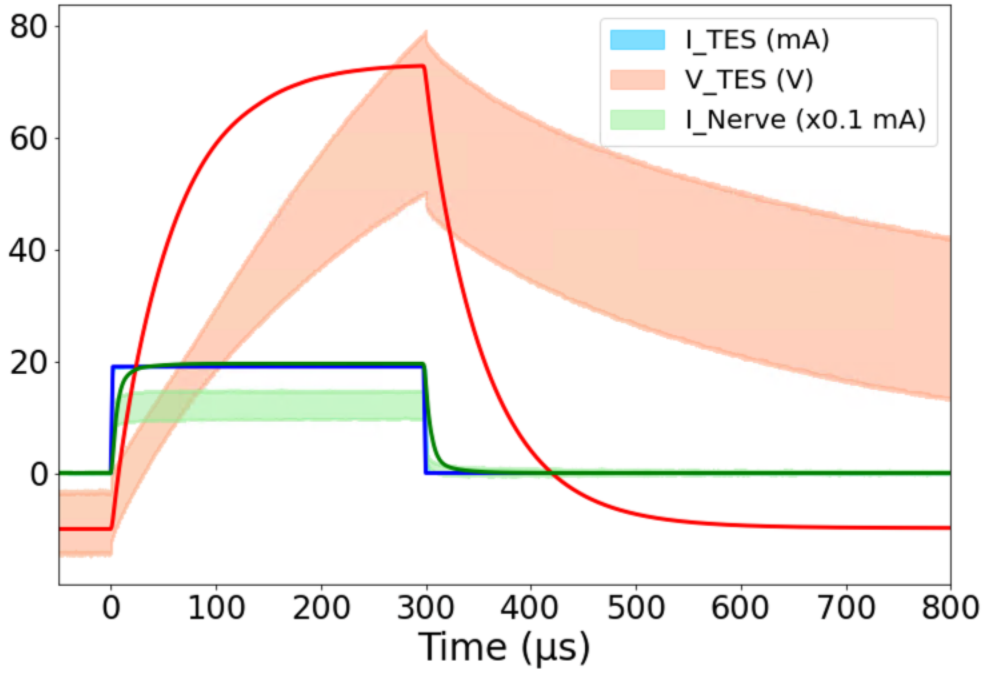


Figure S3a: Current-controlled 300 $\mu$s monophasic pulses. Three solid lines are simulation results, and three shaded areas are cadaver measurements $\pm$ 1 SD (n=8 measurements from both sides of n=4 cadavers). Red solid line (simulation) and shaded area (cadaver validation measurements) represent voltage of applied stimulation waveform, blue represents current through surface electrodes, and green represents nerve current (scaled by x0.1 mA for visualization). The approximately -10 V offset in cadaver TES voltage is due to a direct current (DC) offset from the stimulation system in current-controlled mode. Computational model results for TES voltage were offset by a similar -10 V here for visualization purposes. Voltage-controlled measurements did not face this offset issue.

**Supplementary Material 4 – Cadaver validation fitted with values representative of swine skin**

Here, compared to Fig. 4 in the main manuscript and supplementary material 3, skin conductivity and permittivity values were altered from the Gabriel et al. (1996b) human literature values. Skin conductivity was decreased, and permittivity was increased, matching the swine values more accurately. Pig skin at the abdomen lacks hair follicles and therefore sweat glands – lowering the conductivity when compared to human skin – which has sweat glands even in regions without hair follicles (Avci et al., 2013). With the altered skin conductivity and permittivity values for swine skin, we saw a better fit between the finite element method (FEM) model and swine cadaver measurements.

Human skin conductivity = 1.80 x 10^-4 S/m
Human skin permittivity = 1.17 x 10^3

Fitted swine conductivity = 0.9 x 10^-4 S/m
Fitted swine permittivity = 4.68 x 10^3

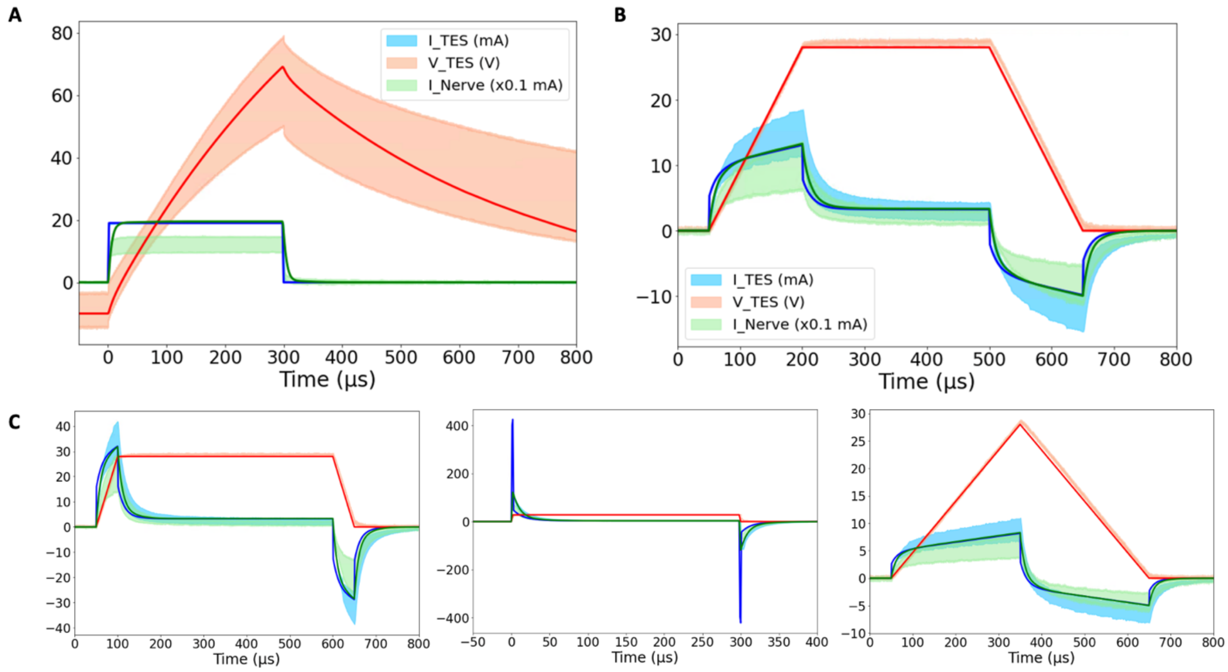


Figure S4a: Domestic swine cadaver verification of FEM model. Three solid lines are simulation results, and three shaded areas are cadaver measurements $\pm$ 1 SD (n=8 measurements from both sides of n=4 cadavers). Red solid line (simulation) and shaded area (cadaver validation measurements) represent voltage of applied stimulation waveform, blue represents current through surface electrodes, and green represents nerve current (scaled by x0.1 mA for visualization). **(a)** Current-controlled 300 $\mu$s monophasic pulses. The approximately -10 V offset in cadaver TES voltage is due to a DC offset from the stimulation system in current-controlled mode. Computational model results for TES voltage were offset by a similar -10 V for visualization purposes. Voltage-controlled measurements did not face this offset issue. **(b)** 28 V voltage-controlled 600 $\mu$s pulses with 150 $\mu$s rise and fall times. **(c)** (Left) 28 V voltage-controlled pulses of 600 $\mu$s duration with 50 $\mu$s rise and fall times. (Center) 300 $\mu$s duration with fastest (~2 $\mu$s) rise and fall time. (Right) 600 $\mu$s duration with 300 $\mu$s rise and fall times. Note: 50 $\mu$s rise time (left) is n=7 measurements due to the incorrect application of waveform amplitude in one sample.

**Supplementary Material 5 – Mechanism of transcutaneous charge transfer**

Despite the exponential capacitive waveforms in Fig. 4 (b-c), the main mechanism of charge transfer to the nerve is ohmic. The TES-tissue interface is highly capacitive, but once charge enters tissue, ohmic charge transfer dominates. This trend was investigated by setting the skin conductivity to 0 S/m while leaving the permittivity unchanged in the transcutaneous coupling FEM model. A transient simulation was run, and charge transferred to the nerve was calculated as area under the rectified $I_{Nerve}$ curve. In the Fig. 4 (c) waveform with the fastest rise time, 29% of the charge transferred to the nerve was maintained when the conductivity of skin was set to 0 S/m and the only way for charge to cross the skin layer was as displacement current.

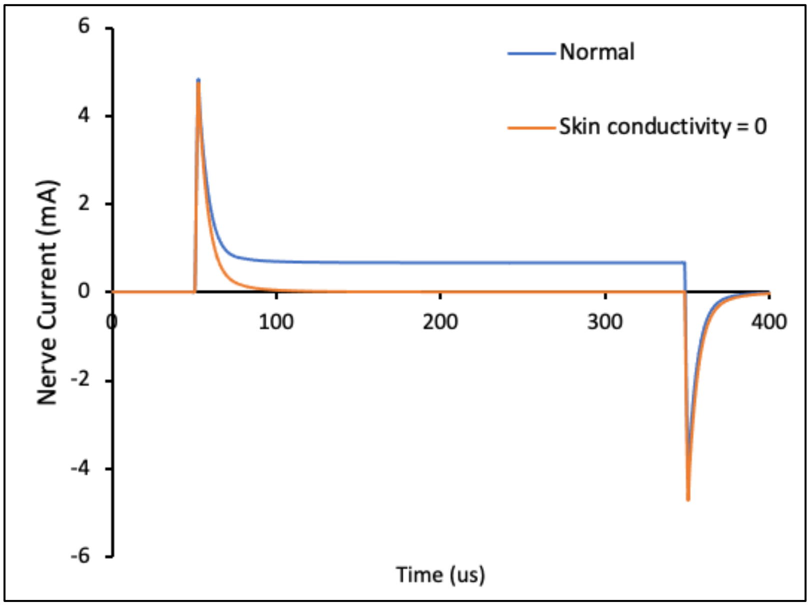


Figure S5a: Nerve current against time during application of a 300 $\mu$s duration voltage-controlled pulse. Blue trace represents normal conductivity and permittivity values of skin. Orange trace represents skin conductivity set to 0 S/m (insulator) and the only mechanism for charge to enter tissue was as displacement current. This allowed us to quantify displacement charge transfer versus ohmic charge transfer into the tissue.
